# The effect of surfactants and film-forming polymers on pulmonary surfactant function measured *in vitro* is dose-rate dependent

**DOI:** 10.1101/2024.10.18.618437

**Authors:** Sreyoshee Sengupta, Hugh J. Barlow, Maria T. Baltazar, Jorid B. Sørli

## Abstract

Surfactants and film-forming polymers are common ingredients in consumer spray products such as cleaning products, hair care products, and anti-perspirants. Spraying eases application by creating aerosolised droplets of the product that can distribute evenly over the treated surface. However, these aerosols can potentially be inhaled during their normal application. Droplets that reach the alveoli can interact with the pulmonary surfactant; a complex mixture of phospholipids and proteins that regulates the surface tension at the air-liquid interface. This interaction could elevate the minimum surface tension at maximum compression and change the surface rheology of the pulmonary surfactant at the interface. We tested four surfactants and seven polymers for their ability to inhibit pulmonary surfactant function *in vitro* and investigated if the inhibition is dose-rate dependent i.e., the product of the concentration (mg/mL) and aerosolisation rate (mL/min). We found that independent of chemical class (surfactant or polymer) there was a clear dose-rate dependent inhibition of pulmonary surfactant function and that different chemicals inhibited function at different dose-rates. We compared the points of departure of inhibitory chemicals to a polymer with known dose-rate dependent lung toxicity. When assessing the risk of chemicals that might be inhaled, it is essential to ensure normal use would not inhibit pulmonary surfactant function leading to immediate effects on the lungs.

**Lay summary:** Spray products create a cloud of tiny droplets in the air when they are used. This cloud can be inhaled, and if it reaches the deepest parts of the lungs, it can interact with the thin layer of liquid, called pulmonary surfactant, that covers the cells. It protects the lung tissue during the constant movement of breathing. Droplets can sometimes disrupt the pulmonary surfactant function, making breathing difficult. Chemicals that are used in spray products must be tested to assess if they are harmful if inhaled. In this project we studied the effect of chemicals that are commonly found in spray products on the functioning of the pulmonary surfactant *in vitro*. The results can be combined with other *in vitro* methods to test if chemicals are harmful to inhale without testing on animals.

## 1. Introduction

In 2010 a large group of people became ill after being exposed to an aerosol of an impregnation product during renovation of a supermarket (Duch et al., 2014). The product was not intended to be sprayed in the manner that it was. The airless spray gun used created small droplets (aerosols) in the air inhaled by workers and customers in the supermarket (Duch et al., 2014). Whereas the accident could have been avoided if the product was not sprayed, many consumer products are formulated as sprays since this form of application ensures easy application and an even distribution of the product over the treated surface area. Aerosols generated during this process can potentially be inhaled by the user or bystanders. Presently, evaluating the risk of chemicals in consumer spray products relies on different strategies, e.g. lung exposure estimation, history of safe use, historical animal data, performing new animal experiments, or testing in so called new approach methods (NAMs), e.g. *in vitro* assays that mimic the lungs. Herein we propose one such NAM as an important assay to test sprayed chemicals, namely testing the effect on pulmonary surfactant function.

Type II alveolar cells produce and release pulmonary surfactant into the liquid lining of the alveoli, where it rapidly spreads and covers the air-liquid interface (Perez-Gil, 2008; Possmayer et al., 2023). The alveolar-capillary membrane, the layer of cells separating the external environment from the blood, is thin and has no structural support for the tissue. By regulating surface tension at the alveolar air-liquid interface, pulmonary surfactant supports the mechanics of the alveoli during breathing. As the surface area of the lungs decreases with an out-breath, the surface tension decreases, thereby preventing the alveoli from collapsing. With the next in-breath, the surface area increases, the phospholipids and proteins rearrange at the air-liquid interface, and the surface tension increases (Perez-Gil, 2008; Possmayer et al., 2023).

Studies performed on the product that caused the supermarket renovation accident, both *in vivo* and *in vitro*, found that the impregnation product was acutely toxic to mice, most likely due to the inhibition of pulmonary surfactant function as observed *in vitro* (Duch et al., 2014; Sørli et al., 2018). The changes that were observed both in exposed people (immediate onset of coughing and dyspnoea) and mice (rapid reduction in tidal volume indicating alveolar collapse), suggest that pulmonary surfactant function inhibition was the mechanism of toxicity (Duch et al., 2014). However, the determination of pulmonary surfactant function was based only on the effect on the minimum surface tension. We hypothesized that the impregnation product modified the rheological properties of the pulmonary surfactant film, and that calculating this change could be used as a tool for characterising hazards of inhalable chemicals. Based on the published literature of case studies involving poisoning by inhalation of impregnation products we also theorised that the toxicity was based on the dose-rate of the exposure, rather than the dose. In this work we use hazard ranking of the substances, as we calculate a point of departure where the rheological properties of the pulmonary surfactant film change from the unexposed state and compare it to a control with a known *in vitro* effect. This information could be used to guide chemical selection, product type and associated exposure, for example, by characterising the particle size distribution of aerosols and sprays to avoid alveolar deposition.

To test these hypotheses, we herein focus on two groups of chemicals; surfactants and film-forming polymers, both are frequently found in different types of consumer spray products e.g. products for cleaning, disinfection, impregnation, and in cosmetics and personal care products. Most of the tested chemicals are already safely used in spray products, however, to our knowledge, one polymer is not presently used in sprays (Polymer 1, see materials and methods). We tested this polymer to determine a hazard assessment compared to the other tested polymers and surfactants.

We have previously shown that different inhaled chemicals can inhibit pulmonary surfactant function *in vitro*, and that this translates to change in lung function in experimental animals (Sørli et al., 2018; Larsen et al., 2020; Hougaard et al., 2023) and in accidentally exposed persons (Sørli et al., 2018; Jensen et al., 2024). The physiological impact on the lungs when pulmonary surfactant function is disrupted is that the surface tension during maximum compression (at the end of an out-breath) is high, and the alveoli collapse. The collapsed alveoli can reopen with an in-breath and collapse again. This puts strain on the alveolar-capillary membrane that separates the air and the blood in the lungs. The process has been described in an adverse outcome pathway (Da Silva et al., 2021c).

Recently, researchers have analysed how the rheological properties of pulmonary surfactant can be modified by interaction with nanomaterials (Sosnowski, 2018; Kondej and Sosnowski, 2019, 2020; Xu et al., 2023) and e-cigarette components (Sosnowski et al., 2018; Xu et al., 2022; Graham et al., 2022; Goros et al., 2023). They demonstrated that the introduction of a non-native chemical species to the pulmonary surfactant system led to a marked change in the interfacial rheology. Furthermore, Zasadzinski and co-authors examined the plausible link between a decrease in the dilational complex modulus and Acute Respiratory Distress Syndrome (ARDS) (Zasadzinski et al., 2010; Sachan and Zasadzinski, 2018; Barman et al., 2020). They suggested that such a decrease may lead to a Laplace instability, causing the alveoli to fail to inflate and deflate during normal breathing. In the present study, we analyse the change in the dilational rheology of pulmonary surfactant due to the introduction of several different aerosolised chemicals. Our study examines model pulmonary surfactant in a constrained drop surfactometer (CDS), exposed to the aerosolised test chemicals at different aerosolisation rates and concentrations, i.e. different dose-rates. The current study aims at quantifying the effect on changes in the viscoelastic properties of the pulmonary surfactant. We have ranked the chemicals in relative potency and used two examples with impregnation products to assess their relative risk.

## 2. Materials and Methods

### 2.1 Materials

We tested four surfactants and six polymers, as well as the impregnation product described in the introduction. Surfactants are amphiphilic molecules with a hydrophobic alkyl chain and a hydrophilic head (Nakama, 2017). Polymers are large molecules composed of repeating units of small molecules called monomers connected by covalent bonds (Ravve, 2012; Lochhead, 2017).

The following chemicals were bought from Sigma-Aldrich (Germany): benzalkonium chloride (BAC), hexadecyltrimethylammonium chloride (HTA), sodium dodecyl sulphate (SDS), Triton X-100 (TX-100), 1H,1H,2H,2H perfluorooctyl triethoxysilane (POTS) and 2-propanol. Ethanol was bought from Histolab products (Denmark). Polyhexamethyleneguanidine phosphate (PHMG) was bought from BOC sciences (USA), and polymethylenebiguanide hydrogen chloride (PHMB) samples were a kind gift from Laboratorie Pareva (France). A sample of Acudyne™DHR (acrylate co-polymer) from Dow Chemical Company was provided by Unilever. Gantrez ES-425 (BE PVM/MA) was provided by Vendico Chemical AB, Sweden. Siloxane polymer (Polymer 1) was provided as a free sample. Stain Repellent Super (SRS) is produced by Akemi GmbH (Germany), it is a mixture of alkyl siloxanes in C9-C13 alkanes (Duch et al., 2014). Abbreviations, CAS numbers, chemical class, harmonized CLP classification, the solvents used for testing, and the chemical structures are listed in table 1. The physical-chemical properties are described in table 2. The physical appearance and stock solutions used can be found in Supplementary table 1.

**Table 1:**
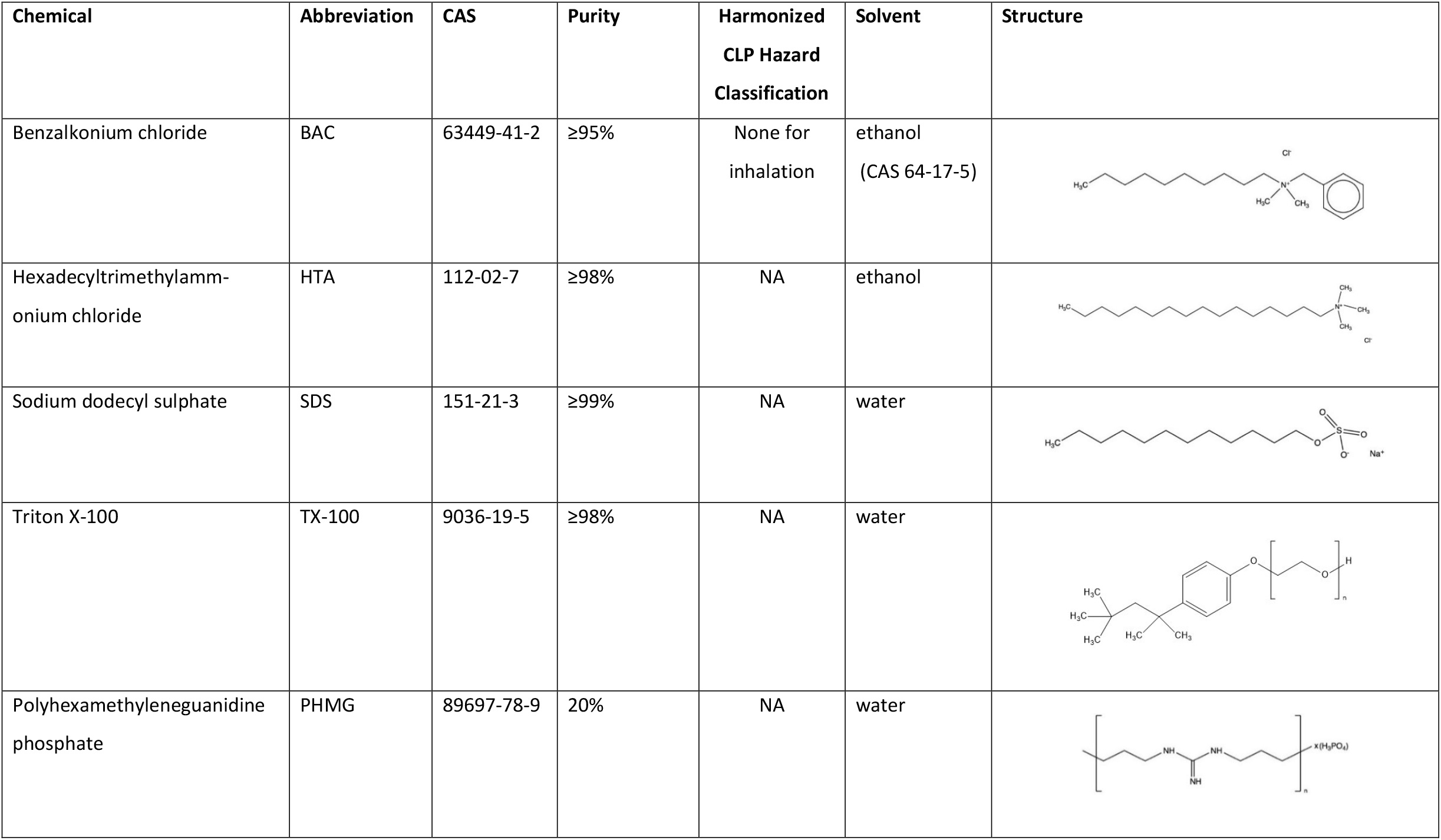
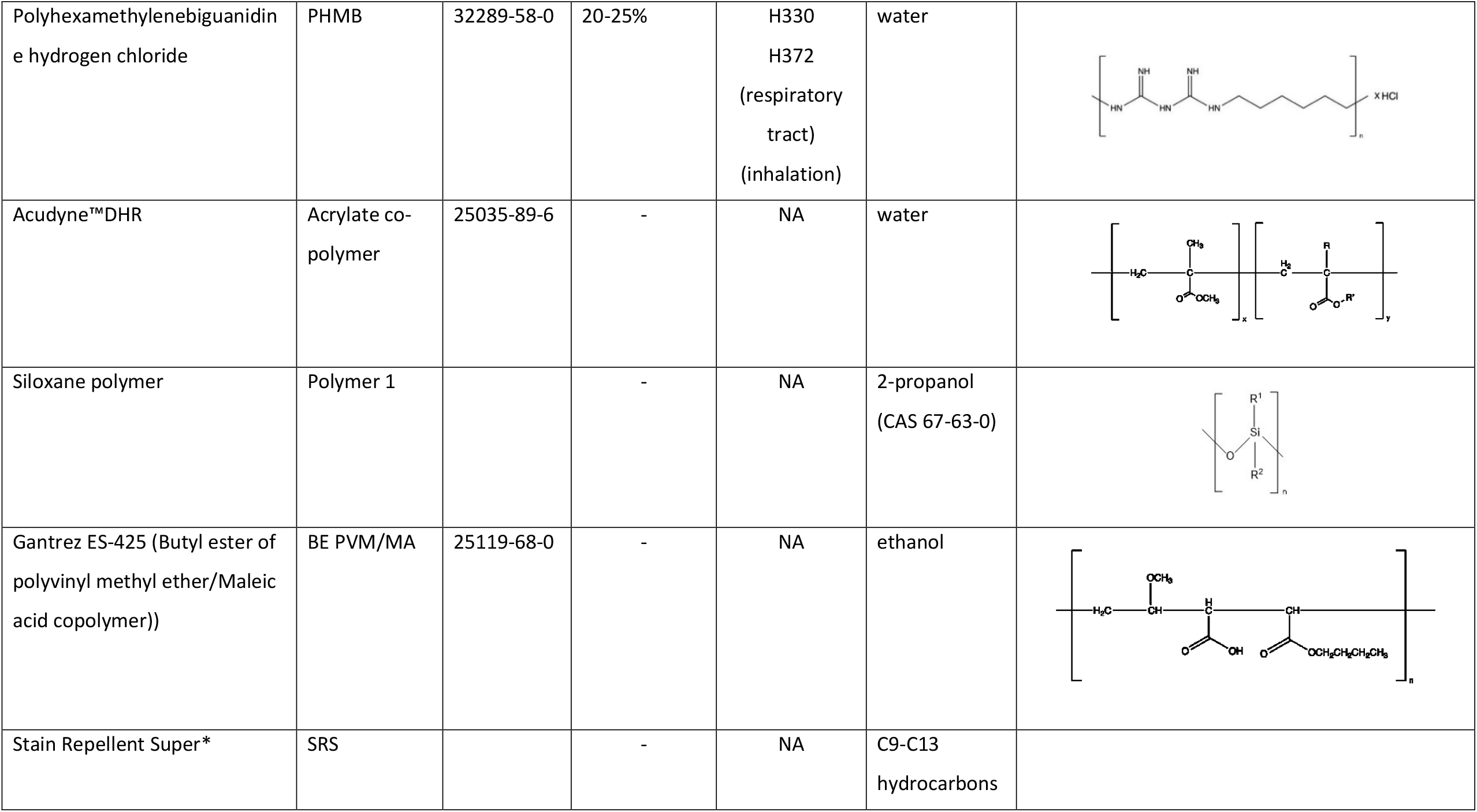
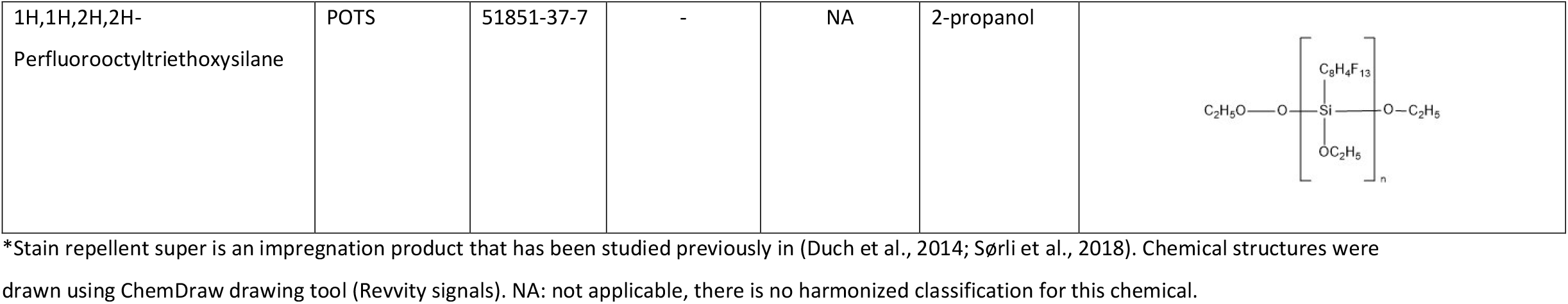
The names and characteristic of the surfactants and polymers tested for pulmonary surfactant function inhibition.

**Table 2:**
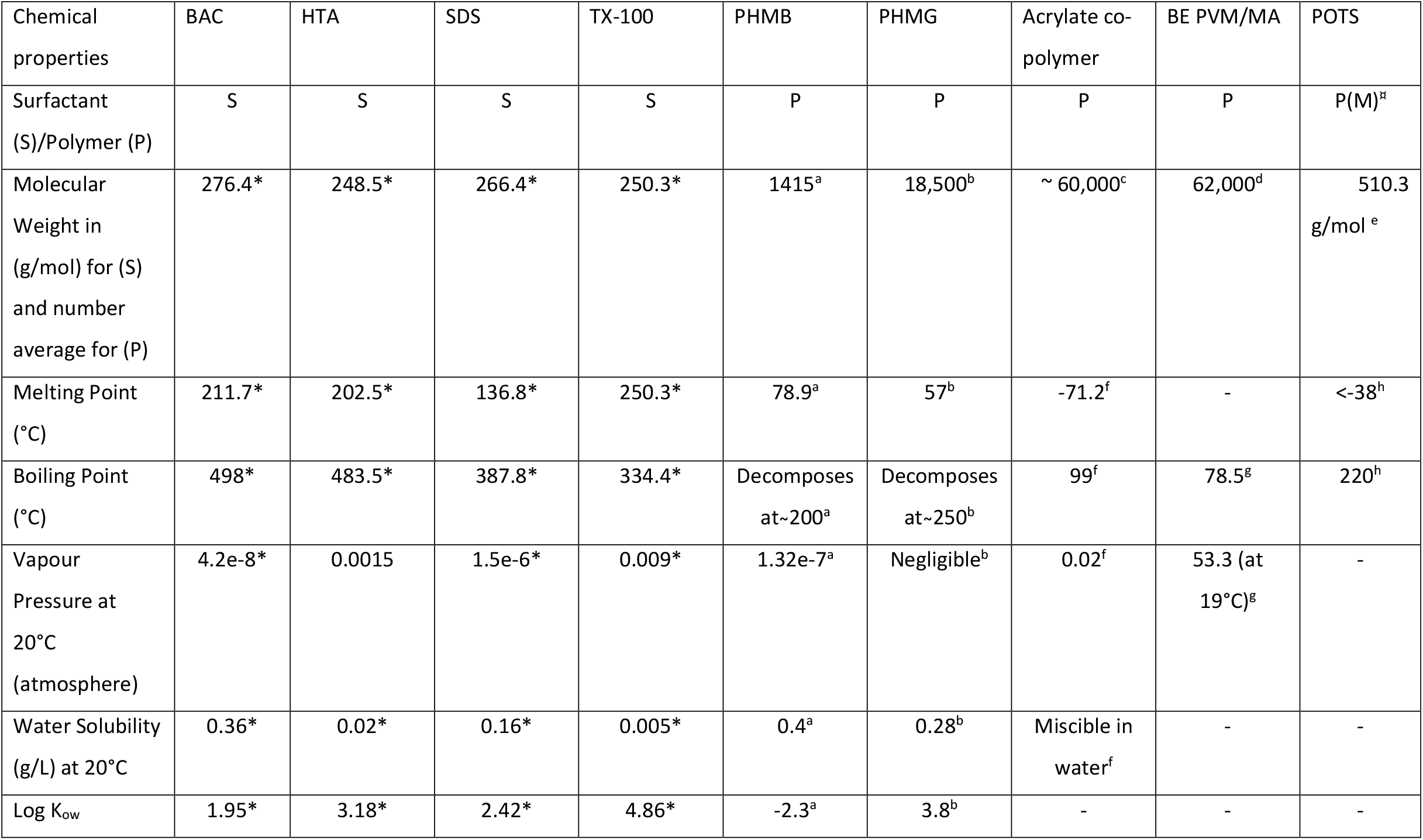
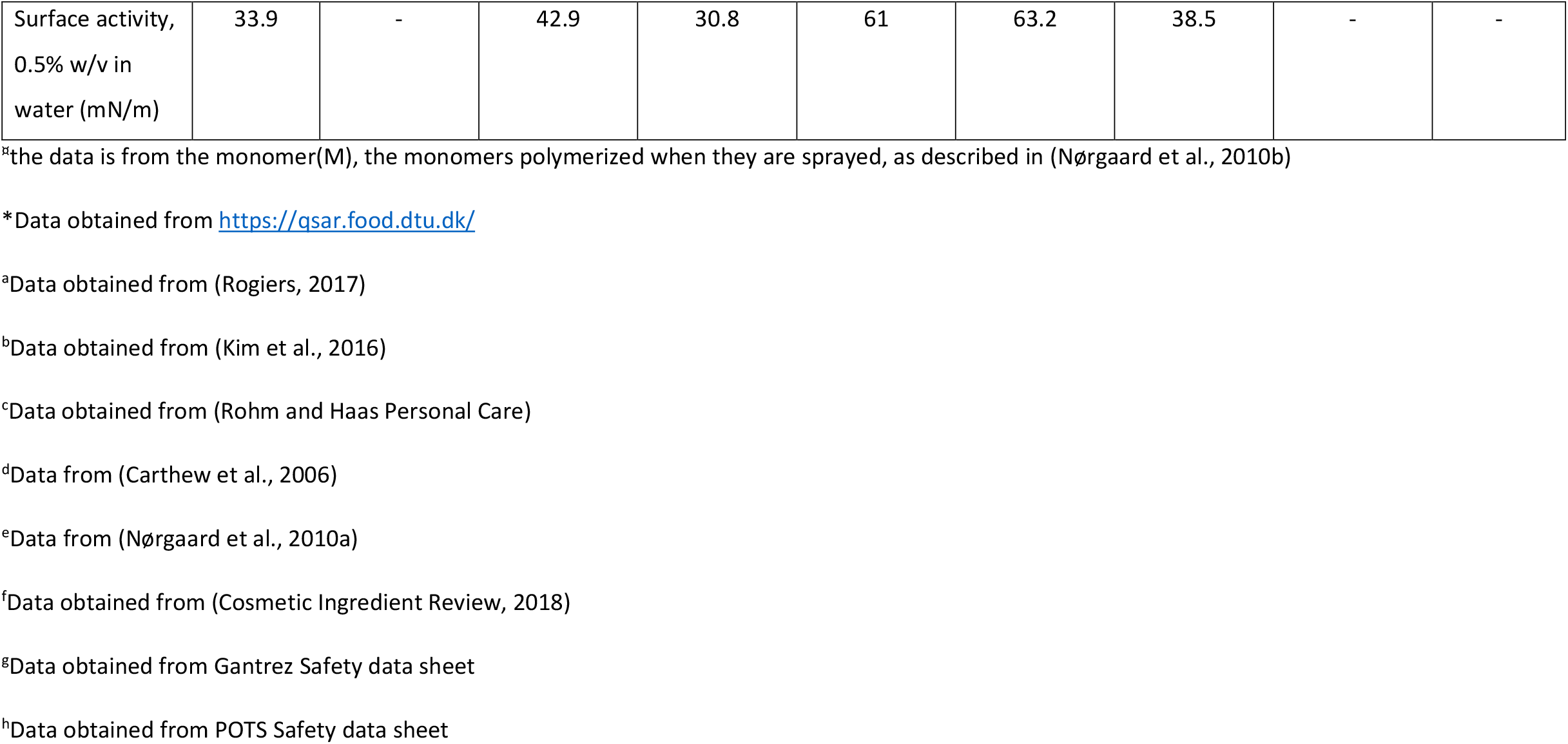
The physical-chemical properties of surfactants and polymers tested for lung pulmonary surfactant function inhibition.

The pharmaceutical Curosurf^®^ (Chiesi, Italy) was used as a model for pulmonary surfactant. Curosurf^®^ is an animal-derived product, made from solvent extracted minced pig lung tissue and contains ∼99% w/w phospholipids and 1% w/w hydrophobic surfactant associated proteins (SP-B and SP-C) (Possmayer et al., 2023).

### 2.2 Surfactant preparation

Curosurf^®^ is supplied as a pharmaceutical 1.5 mL dose with a phospholipid concentration of 80 mg/mL. The vial was aliquoted, freeze dried, and stored at -80°C so that a fresh batch could be used every experimental day. On the day of experimentation, the aliquot was thawed and reconstituted using a buffer containing 0.9% NaCl, 1.5 mM CaCl_2_, and 2.5 mM HEPES, adjusted to pH 7.0 (Sørli et al., 2022) to a concentration of 2.5 mg/mL.

### 2.3 Constrained drop surfactometer experiments

The Constrained Drop Surfactometer (CDS, BioSurface Instruments, USA) was used to measure the surface tension of pulmonary surfactant under dynamic conditions (Sørli et al., 2016; Xu et al., 2023). The CDS setup was used with a constant flow through exposure chamber as described in (Sørli et al., 2016) to study the effect of aerosolised test chemicals on pulmonary surfactant function.

A drop of pulmonary surfactant (Curosurf, 10 µL at 2.5 mg/mL) was placed on a hollow, sharp edged pedestal connected via plastic tubing to a syringe filled with buffer, the syringe was placed in a computer controlled motorised pump. The drop was subjected to dynamic cycling (20 cycles/minute, at 20-30% compression) by introducing and removing liquid from the drop through the base of the pedestal to simulate breathing. The cycling of the drop was stopped to refill the drop with buffer and replace the evaporated liquid when the drop became too small. Pulmonary surfactant was cycled for 40 seconds to obtain a baseline value followed by exposure to the aerosolised chemicals for 5 minutes. Experiments with a baseline minimum surface tension value >5 mN/m or compression >30% were discarded. Drop images were continuously recorded at five frames per second, and analysed by axisymmetric drop shape analysis (ADSA) software (Yu et al., 2016) to calculate surface tension and surface area among other parameters. The temperature in the exposure chamber was monitored using the TinyTag Plus 2 data logger (TGP-4017, Gemini Data Loggers Ltd, United Kingdom). The CDS setup was placed in a heating box and the air used to aerosolise the test chemical was heated. Aerosols of the test chemical were generated by infusing a solution of chemical in solvent (Table 1) with an infusion pump (Legato 100, Buch & Holm A/S, Denmark) into a Pitt no.1 nebulizer (Wong and Alarie, 1982) by the flow of warm pressurized air into the nebulizer, i.e. the aerosolisation rate.

For each test chemical, the pulmonary surfactant was exposed to different concentrations of the chemical, and each concentration was tested at three different aerosolisation rates (0.1 mL/min, 0.25 mL/min and 0.5 mL/min). Each combination of concentration and aerosolisation rate was repeated 4-6 times for each chemical. After each experiment the pulmonary surfactant was removed with a tissue, buffer was pushed through the pedestal holder 3-4 times, removing the buffer with a tissue soaked in ethanol.

#### 2.3.1 Positive control (POTS)

A positive control, 1.3% 1H,1H,2H,2H-perfluorooctyltriethoxysilane (POTS) in 2-propanol was included on each experimental day at an aerosolisation rate of 0.1 mL/min to check the performance of the system. The chemical was previously studied in(Nørgaard et al., 2010a; Larsen et al., 2014; Nørgaard et al., 2014; Sørli et al., 2018; Sørli et al., 2022), and the data (among other) was used to generate a restriction in REACH (European Commision, 2019) that prohibits POTS in organic solvents from being sold as a spray product. Exposure to this product inhibited pulmonary surfactant function *in vitro* and caused an irreversible reduction in tidal volume leading to lung damage in exposed mice (Nørgaard et al., 2010a; Sørli et al., 2018). We followed the instructions in to produce a POTS solution with known effect on the respiration of exposed mice(Nørgaard et al., 2010a). Thus POTS was prepared by mixing 54 µL of water (Mw 18 g/mol, density 1 g/mL), 11 µL formic acid ≥95% (Mw 46 g/mol, density 1.22 g/mL), 392 µL 1H,1H,2H,2H-perfluorooctyltriethoxysilane (POTS) 100% (Mw 510 g/mol, density 1.3 g/mL), and 100 µL 2-propanol 99.9% (Mw 60 g/mol, density 0.8 g/mL) in a 2 mL plastic Eppendorf tube. The vial was clamped in an overhead shaker (REAX 2, Germany) and gently mixed at a speed of 20 rounds per minute (rpm) for 2 hours to obtain a homogenised, translucent 77.6% wt/wt POTS chemical mixture. The mixture was added to 36.6 mL of 2-propanol in a low-density polyethylene flask and mixed to obtain a 1.3% wt/wt, or 13.7 mg/mL POTS solution. The mixture was stored at room temperature. POTS was tested as the other test chemicals, i.e. by altering the concentration and the aerosolisation rate (Fig. 1).

**Figure 1:**
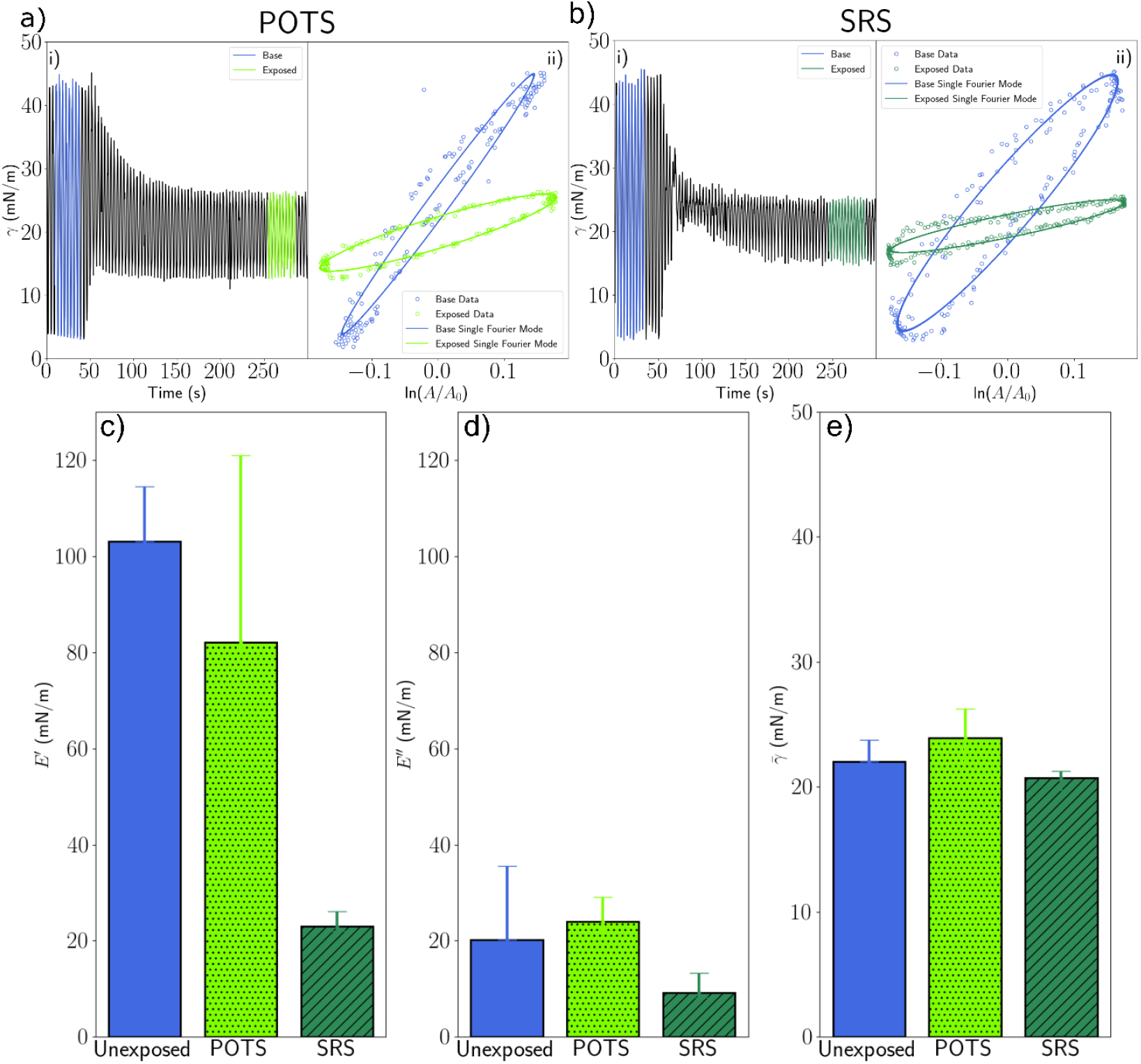
Surface dilational rheological changes when pulmonary surfactant interacts with aerosolised POTS or SRS. a)i) and b)i) An increase in minimum surface tension of pulmonary surfactant during compression is observed after the pulmonary surfactant is exposed to aerosolised POTS (concentration of POTS 13.7 mg/mL) and SRS both at aerosolisation rate of 0.1 mL/min. The pulmonary surfactant drop was cycled for 40 seconds (baseline) before aerosolisation of the test chemical was started. a)ii) and b)ii) Hysteresis curves based of largest Fourier mode (lines) obtained from the subset data (dots) chosen before (blue) and after exposure (green). (c), (d) Viscoelastic properties such as dilational elastic modulus E’, and dilational viscosity modulus E’’ respectively obtained for the subset before and after exposure; a drastic decrease in the dilational elastic modulus and small change in the dilational viscous modulus is observed and (e) average surface tension 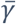 where almost no change is observed before and after exposure.

#### 2.3.2 Negative control

Each experiment has a baseline period with unexposed pulmonary surfactant. Chemicals that were tested in a water solution did not have a negative (solvent) control for each experimental day. Chemicals dissolved in other solvents were tested with the respective solvents as a negative control on each experimental day (i.e. ethanol or 2-propanol).

#### 2.3.3 Surface activity

To determine if the chemical had surface activity (inspired by the ASTM standard D1331, but by using the CDS), it was dissolved at a concentration of 0.5% in water. A drop of the solution (30 µL) was placed on a 5 mm diameter pedestal and allowed to stabilise for 5 minutes, followed by 1 min where pictures of the drop were taken at one frame per second. The mean surface tension during these 60 seconds is summarised in table 2, the temperature in the chamber was at room temperature. Surface active chemicals were defined as having a surface tension of ≤45 mN/m during the 1 min of data collection. Chemicals that were not possible to dissolve in water (HTA, polymer 1 and BE PVM/MA and POTS) were not tested for surfactant criteria since the solvents are volatile and have a very low surface tension in themselves.

### 2.4 Literature review

To identify if there was literature on the effects of the chemicals on pulmonary surfactant, or other parts of the lungs we searched PubMed for relevant literature. The literature search strategy and results are summarised in supplementary table 2.

### 2.5 Rheology

#### 2.5.1 Surface dilational rheology of pulmonary surfactant

The surface tension of a pulmonary surfactant monolayer exhibits complex behaviour as a result of the dilation undergone during breathing (Wustneck et al., 2005; Ravera et al., 2021). This is due to the surface active phospholipids (enriched in dipalmitoylphsophatidylcholine (DPPC)) and proteins that adsorb/desorb, respread and rearrange at the air-liquid interface during the breathing cycles. Compression and expansion cycles result in periodic oscillations in the surface tension. Furthermore, the surfactant film forms biologically complex interfaces comprising of semi-crystalline liquid condensed and disordered liquid expanded phases respectively (Possmayer et al., 2023). This result in changes in the shear and dilational deformations of the interface. Dilational rheology of the pulmonary surfactant monolayer characterises the dynamic surface tension in response to the change in surface area of this system (Wustneck, 2001). For a purely elastic system, the dilational modulus (E) is defined as change in surface tension divided by the logarithmic change in area (Equation 1)

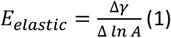

where *γ* and *A* are the surface tension and surface area respectively. Due to the kinetics of molecular motion between the surface of the droplet and the bulk pulmonary surfactant being on timescale comparable to breathing frequency, 12-18 breaths per minute, there is also a viscous response to deformation, characterised by the dilational viscosity

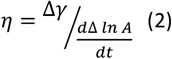

In a periodically oscillating system, the elastic and viscous responses are combined as the dilational complex modulus E^*^ that consists of the elastic component, the dilational elastic modulus (E’) and the viscous component, the dilational viscous modulus (E’’). The complex dilational modulus at a given frequency of oscillation *ω* is given by

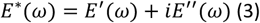

In this study, the viscoelastic properties were investigated by studying the surface dilational rheological parameters of the surfactant monolayer at a fixed frequency of 20 breaths per minute. Previous studies of the pulmonary surfactant have stressed the importance of these dilational parameters to the function of breathing (Wustneck, 2001; Alonso et al., 2004; Wustneck et al., 2005; Sachan and Zasadzinski, 2018; Bykov et al., 2019). High surface elasticity rather than very low minimum surface tension was considered as the deciding factor for normal lung function in (Lunkenheimer et al., 1996).

#### 2.5.2 Analysis

In this study, we investigated the changes in the viscoelastic properties of pulmonary surfactant when exposed to an aerosolised chemical during dynamic cycling in the CDS. The dilational complex modulus was determined from the time course data using the equation

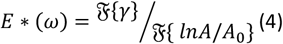

where ℱ denotes the Discrete Fourier Transform operator, *γ* the surface tension, A is the area of the droplet and *A*_0_ is the initial droplet area. Similar processes have been used in the past to study the dilational rheology of surfactant monolayers and are described in detail in Refs. (Ravera et al., 2005; Vrânceanu et al., 2007; Ravera et al., 2010; Yang et al., 2019).

In Figure 1, we see how this method is applied to our datasets. A subset of the data before and after the introduction of the aerosol is selected and the dilational viscoelastic properties are measured using Equation 3. For a pulmonary surfactant droplet exposed to POTS or the impregnation product SRS, the aerosolisation data and the corresponding hysteresis curves of the selected subsets are shown in Figure 1 a), and b) respectively. Also shown is the theoretical hysteresis curve which corresponds to only the largest Fourier mode of the data i.e. the frequency of the applied oscillation. We see that despite some non-linearity in the response to the oscillation of the droplet, the theoretical curves approximate the data very well, thus demonstrating that the dilational properties of the pulmonary surfactant can be quantified using just the complex dilational modulus *E*^∗^ and the average surface tension 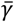.

In Figure 1 c), d) and e), we see the magnitude of these properties before and after the model pulmonary surfactant is exposed to POTS or SRS. We note that there is a dramatic drop in the elastic modulus and a small change in the viscous modulus, but almost no change in the average surface tension. This study will examine how these properties change across a suite of commonly aerosolised chemicals. To quantify any measured change in the viscoelastic response, we define the normalised difference in the complex dilational modulus 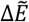 as

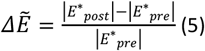

where *E*^∗^_*p*re_ and *E*^∗^_*post*_ are the dilational complex moduli before and after the exposure of the droplet to the chemical and 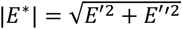 is the magnitude of complex modulus *E*. The corresponding change in the average surface tension 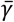 is quantified by the expression

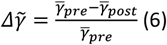

Where 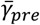 and 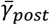 are the average surface tension before and after exposure is initiated. In both cases, we normalise the change to minimise the impact of sample variability in the pulmonary surfactant, or the influence of external uncontrollable factors on the base behaviour.

#### 2.5.3 Estimating a Point of Departure

For each chemical studied here we determine the point of departure by fitting to a mechanistic model derived in (Barlow et al 2024, DOI of preprint on bioRxiv: 10.1101/2024.10.16.618442). We briefly outline the model here without discussing it in detail and then describe how it was used to derive a point of departure.

The model is based on the kinetics of adsorption and desorption of a surfactant system. The evolution of the surface tension as a function of the concentration per unit area of pulmonary surfactant *Γ* and test chemical *Γ*_*x*_ is modelled using the equations

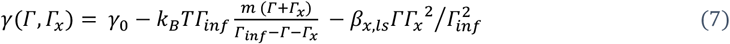

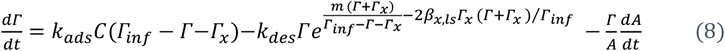

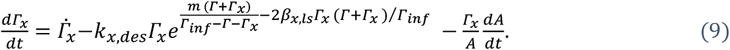

Here *γ*_0_ is the surface tension of water at zero surfactant concentration (set at 71.2 mN/m), T is the temperature (assumed 37 °*C*), *k*_*B*_ is Boltzmann’s constant, and m is an empirical scaling parameter which accounts for the additional elasticity due to interactions between surfactant proteins and phospholipids. *Γ*_*inf*_ is the surface concentration at saturation, which is set from the findings of previous studies to 0.02Å^−2^(Liekkinen et al., 2020; Bouchoris and Bontozoglou, 2021; Stachowicz-Kusnierz et al., 2022). The parameters *k*_*ads*_*C* and *k*_*des*_ are the adsorption and desorption constants of the pulmonary surfactant molecules from the droplet surface. *A* is the surface area of the droplet. The parameters 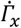, *k*_*x,des*_, and *β*_*x,ls*_ relate to the introduced test chemical, where 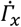 is the rate of deposition of the test chemical per unit area of the droplet, *k*_*x,des*_ is the desorption rate constant of the test chemical and *β*_*x,ls*_ is a parameter which describes the interaction between test chemical molecules on the surface of the droplet. There are multiple stages to fitting this model. First is to establish a base state. The model with 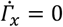, is fit to data from unexposed model pulmonary surfactant by optimizing the parameters *k*_*ads*_*C, k*_*des*_ and *m* to minimise the errors between the model and the data. The base model parameters are included in Table 3.

**Table 3:** Parameters of the base model derived from the mechanistic model of unexposed pulmonary surfactant.

| Parameter | m | $k_{ads}C$ | $k_{des}$ | $\Gamma_{inf}$ | $\gamma_0$ |
| --- | --- | --- | --- | --- | --- |
| Units | # | $s^{-1}$ | $s^{-1}$ | $\text{\AA}^{-2}$ | mN/m |
| Value | 3.11 | 0.045 | 2.09e-05 | 0.02 | 71.2 |

Fixing these parameters, we now model the effects of the test chemical. As we are unable to obtain the exact dose-rate in this study due to experimental limitations, we introduce a new parameter *α* such that 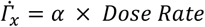. For each test chemical, we therefore fit the model with three parameters, *k*_*x,des*_, *β*_*x,x*_, and *α* to the quantities 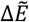 and 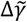 as a function of dose-rate using the scipy optimize library.

## 3. Results

Experiments with surfactants and polymers were performed at multiple aerosolisation rates and concentrations and 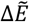 was measured for each of these experiments over multiple repeats (n = 4-6 per test chemical). This study hypothesises that the inhibition of pulmonary surfactant function is *dose-rate* dependent i.e. the product of the concentration and aerosolisation rate which determines the adverse effects on pulmonary surfactant function. The heat maps showing the effect of exposure on the 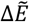 of pulmonary surfactant from exposure to SDS (a surfactant), PHMB (a polymer) and POTS (the positive control) are plotted in Figure 2. We observe that as both concentration and aerosolisation rate were increased, 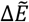 increases, with no difference in trend between increasing concentration or aerosolisation rate. We next plot the values in these heatmaps against dose-rate (Fig. 2). The points plotted display a reasonable curve collapse with several points measured at different aerosolisation rate/concentration combinations showing overlap or seem to fall along the same trend. We also see that the trend is similar in all three test chemicals, a gradual increase in 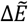 with increasing dose-rate, which seems to flatten as the dose-rate increases. In all cases, the model fits the data very well, suggesting that it may accurately describe the mechanism for inhibition.

**Figure 2:**
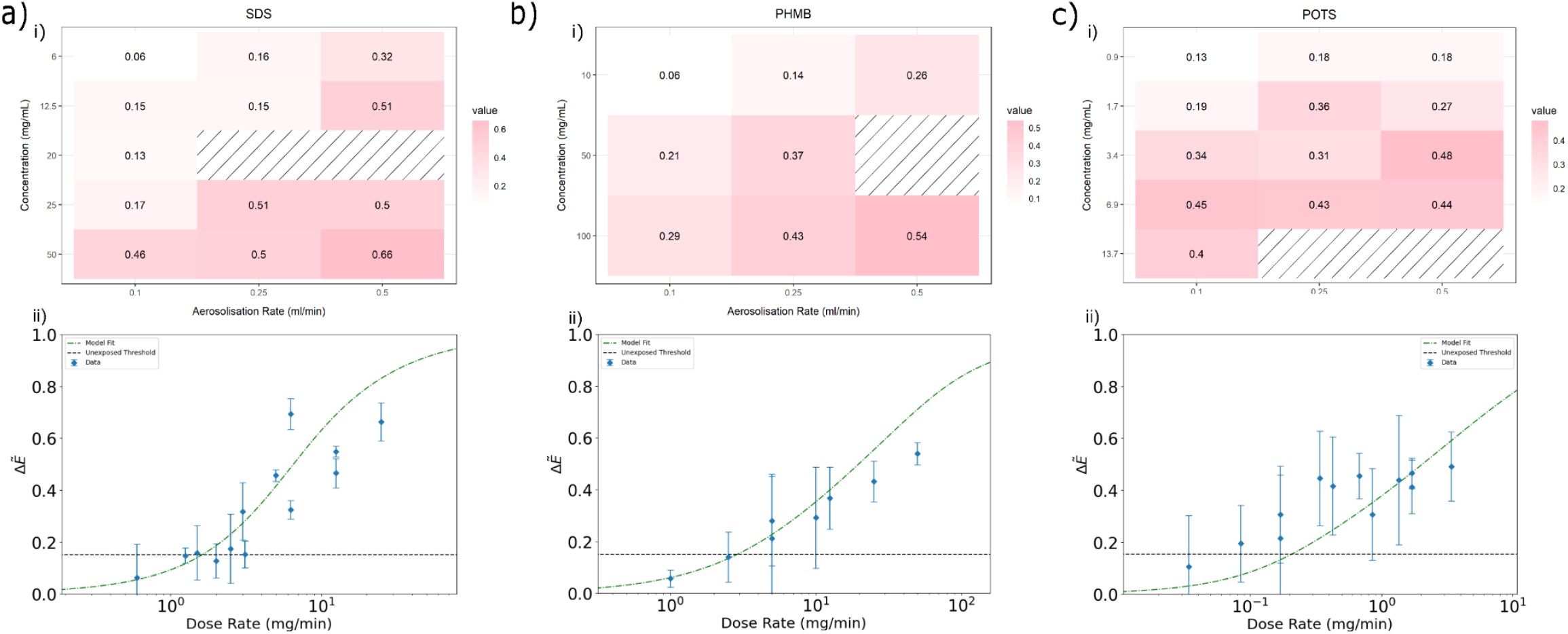
Effect of exposure of SDS (surfactant) and PHMB (polymer), POTS (positive control) on pulmonary surfactant is dose-rate dependent. Heatmaps in figures a, b, c) i) shows an increase in the normalised difference of the complex moduli 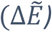 with an increase in both concentration (mg/mL) and aerosolisation rate (mL/min). Corresponding curve collapses a, b, c) ii) are obtained by plotting 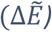 values as a function of dose-rate (the product of concentration and aerosolisation rate) shows a gradual in increase in 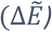 values with a gradual increase in dose-rate. The black broken line shows the unexposed threshold which was used to obtain the point of departures (POD) of SDS (1.63 mg/min), PHMB (2.90 mg/min), and POTS (0.06 mg/min).

The fitted parameters are shown in Table 4 and the fits are shown as broken lines in Figures 2 and 3. The point of departure (POD) for all test chemicals was determined by performing a number of experiments where no test chemical was infused into the chamber. The viscoelasticity before and after 40 seconds was measured and the 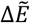 calculated. The average and standard deviation were measured and used to calculate the 95% confidence level, shown as a broken black line in Figures 2, 3, and 4. Using the model described above, the POD was determined as the dose-rate at which 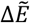 would exceed this level. It is listed along with the model parameters in Table 4.

**Table 4:** Fitted parameters for the tested surfactants and polymers.

| Chemical | $k_{x,des}$ | $\beta_{x,ls}$ | $\alpha$ | POD |
| --- | --- | --- | --- | --- |
| | $s^{-1}$ | $k_B T \Gamma_{\infty}^{-1}$ | $\text{\AA}^{-2} \text{mg}^{-1}$ | mg/min |
| PHMB | 1.66e-05 | -12.178 | 7.90e-5 | 2.90 |
| SDS | 0.0054 | -6.477 | 0.00186 | 1.63 |
| BE PVM/MA | 7.22e-5 | -9.1278 | 0.00023 | 0.74 |
| HTA | 0.0378 | -5.726 | 0.033285 | 0.21 |
| Polymer 1 | 0.00095 | -2.32 | 0.0029 | 0.20 |
| POTS | 9.17e-05 | -5.0255 | 0.0189 | 0.06 |
| BAC | 0.00023 | -6.0 | 0.00761 | 0.04 |

**Figure 3:**
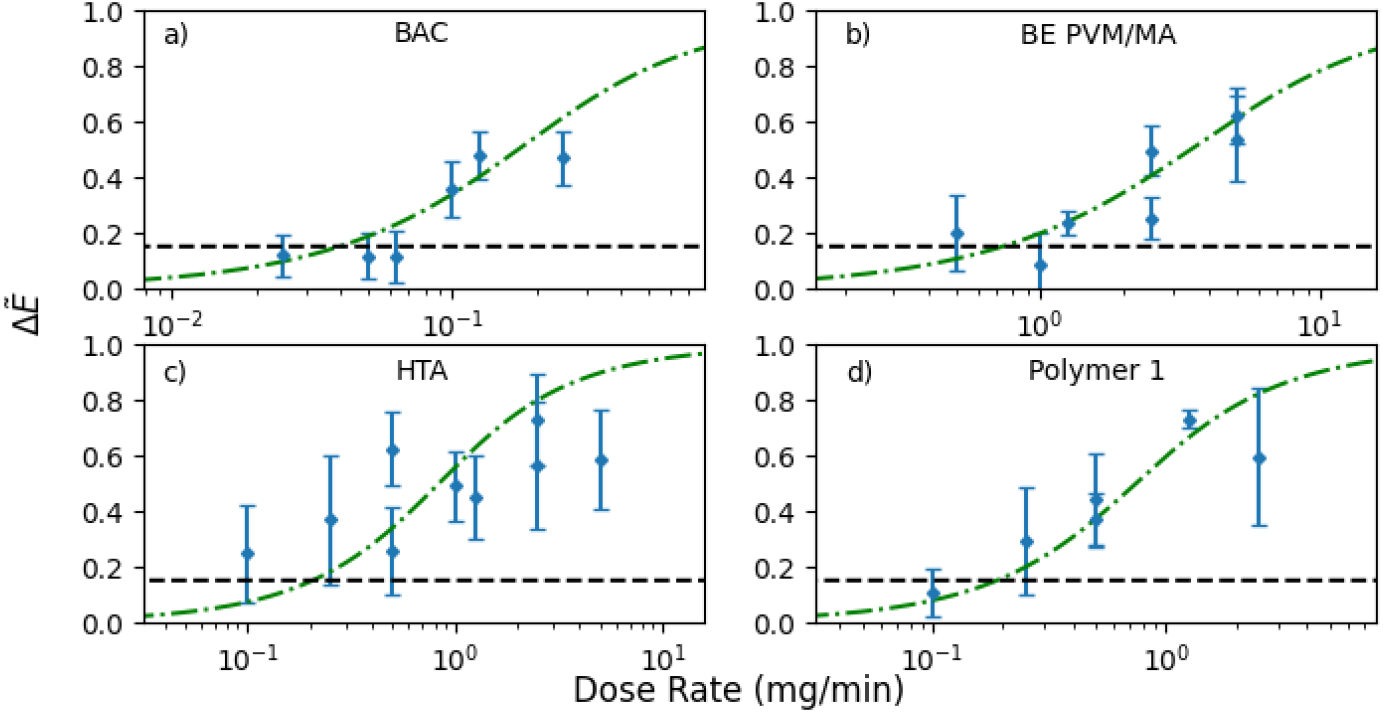
Establishing the generality of dose-dependent hypothesis for both surfactant and polymer classes of chemicals: (a) and (c) are surfactants and (b) and (d) are polymers that inhibit pulmonary surfactant function as the dose-rate is gradually increased. The black broken line shows the unexposed threshold which was used to obtain the POD of BAC (0.04 mg/min), BE PVM/MA (0.74 mg/min), HTA (0.21 mg/min) and Polymer 1 (0.19 mg/min).

**Figure 4:**
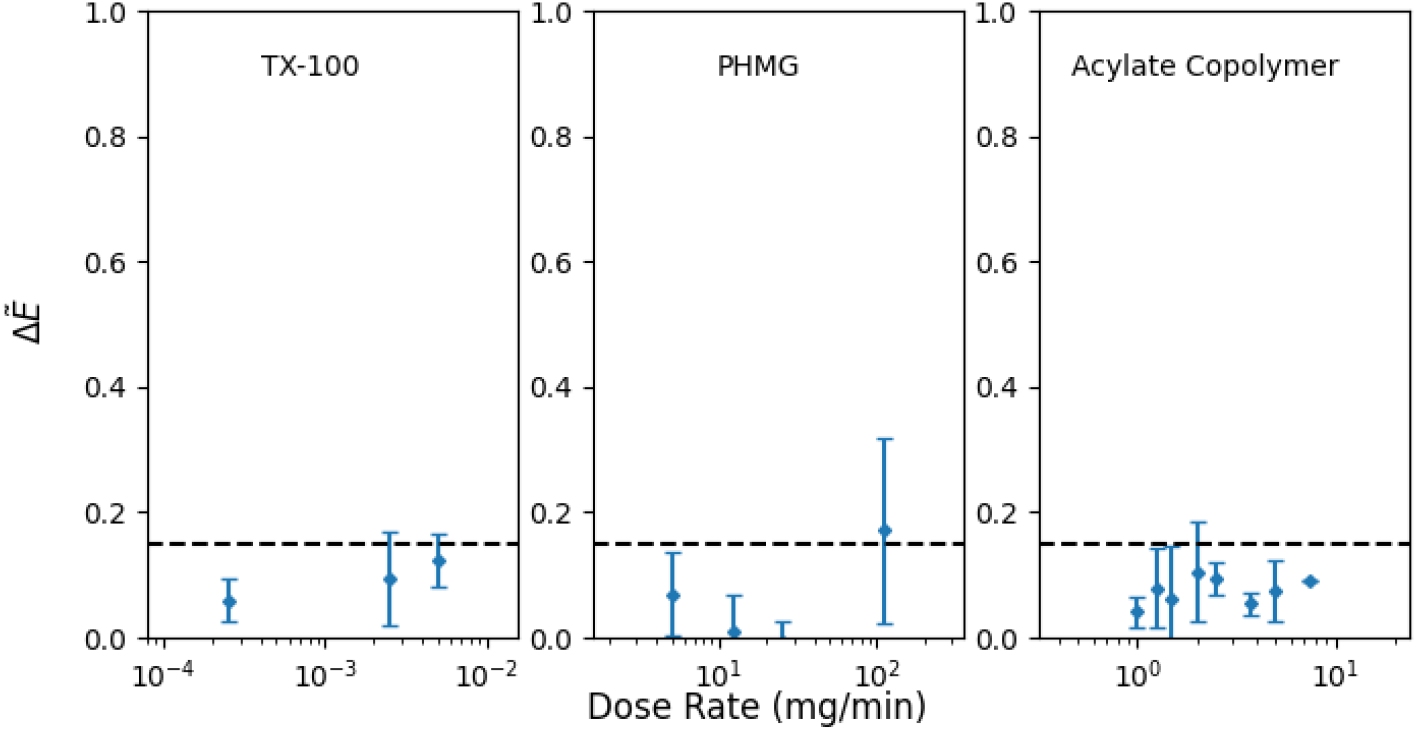
Pulmonary surfactant function inhibition was not observed in Triton X-100 (surfactant), PHMG, and acrylate co-polymer (polymers). 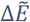 values do not increase with the increase in dose-rate.

Seeking to establish the generality of our hypothesis, we now look at the other surfactants and polymers. Four of these were shown to cause inhibition (Figure 3; BAC, BE PVM/MA, HTA, and polymer 1) and induced the same effects as we saw for SDS, PHMB and POTS. 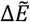 is plotted as a function of dose-rate for each of these test chemicals, and each seems to follow the same trend as seen previously, once again substantiating our claim that the impact of aerosols on pulmonary surfactant function is dose-rate dependent. The kinetics-based model fits the data very well, again suggesting that we have successfully shown that the mechanism for inhibition is generic.

In the case of three test chemicals (Figure 4; TX-100, PHMG and Acrylate co-polymer), no inhibition was observed at the concentrations and aerosolisation rates studied. It is surprising that we do not see any effects for two of these chemicals, TX-100 and PHMG, since they are widely used as anti-microbials and known membrane disruptors. However, effects may be observed at higher concentrations or aerosolisation rates not accessible to this study.

In all cases where inhibition was observed, we see a monotonic increase in the rheological properties which increases with dose-rate. Despite variations in concentration and aerosolisation rate, we see that a dose-rate dependence appears to be universal across the studied chemical space. Where we do see outliers, this may be the result of higher concentration samples forming differently sized aerosols due to changes in viscosity of the fluid being nebulised. Future work examining the exact deposited mass would be able to discern this and also be able to verify the mechanism postulated by our kinetics-based model.

## 4. Discussion

We hypothesised that aerosolised chemicals could change the rheological properties of the pulmonary surfactant film when exposed *in vitro* and that this inhibition was dose-rate dependent. To test these hypotheses we tested an impregnation product SRS, “stain repellent super” from Akemi, that caused 39 people to become acutely ill after accidental exposure (Duch et al., 2014) as well as ten other single chemicals, either surfactants or polymers. We used a chemical (POTS) with a restriction from being sold as a spray product because it inhibits pulmonary surfactant function *in vitro* and causes irreversible lung damage *in vivo* (Sørli et al., 2018) to hazard rank the tested substances. There was an indication that the toxic effects of SRS to humans were dose-rate dependent as the three individuals that were most severely affected were either exposed for a short time to a high concentration (the operator of the spray gun) or for a longer time at a lower concentration (working in the treated area after spraying). Furthermore, when SRS was tested on mice, reduction in tidal volume and concomitant increase in respiratory rate was also dose-rate dependent (Duch et al., 2014). SRS was later found to inhibit pulmonary surfactant function when aerosolised in the CDS (Sørli et al., 2018).

When we aerosolised SRS, the exposure changed the rheological properties of pulmonary surfactant at the lowest possible exposure rate (Fig. 1). Inhibition occurred immediately after the start of exposure which is seen as an increase in minimum surface tension and a simultaneous decrease in maximum surface tension (Fig 1a). Since a change was observed at the lowest aerosolisation rate, the effect of dose-rate *in vitro* could not be determined. The film forming polymer in SRS is alkylsiloxanes in C9-C13 hydrocarbons. As the complete chemical composition of the product is not known, it was not possible to change the solvent, thus the POD of SRS is <77 mg/min, based on the density of the complete product. The POD is likely much less as at this dose-rate the change in viscoelastic properties of pulmonary surfactant was immediate and severe (Fig 1). We could not determine the effects of the single constituents in the SRS product, however, to test our hypothesis, we changed the dose-rate of the tested surfactants and polymers by changing either the aerosolisation rate or the concentration in the feed solution.

### 4.1 Risk assessment of spray products and constituents

For any novel ingredient and product format, a risk assessment is required to assure the safety for consumers during its use. Therefore, comprehensive exposure (Steiling et al., 2014) and hazard assessment is conducted (Scientific Committee on Consumer Safety (SCCS), 15 May 2023; United States Environmental Protection Agency). Alternatives to animal testing have been developed to address this complex endpoint, from exposure based waiving approaches (Carthew et al., 2009) to complex 3D model tissues representing the different regions of the lung (Li et al., 2018; Cao et al., 2021; Ramanarayanan et al., 2022; Mallek et al., 2023). Addressing such diversity of possible adverse effects caused by inhaled chemicals, likely needs a battery of NAMs to cover all relevant molecular initiating events and key events (Clippinger et al., 2018a; Clippinger et al., 2018b). Previous work has demonstrated that measuring pulmonary surfactant function *in vitro* was able to identify liabilities due to interaction with pulmonary surfactant (Larsen et al., 2020; Sørli et al., 2020; Da Silva et al., 2021b; Da Silva et al., 2021c; Jensen et al., 2024), and therefore we hypothesised that this NAM could be part of Integrated Approaches to Testing and Assessment (IATA) for consumer spray products, including cosmetic products. Surfactants and polymers were chosen because 1) they are frequently found in spray products and therefore are potentially inhaled, 2) pulmonary surfactant function is based on the alteration of the surface tension of the air-liquid interface in the alveoli during breathing and as surfactants are also amphipathic it is likely that they would interact with the air-liquid interface, 3) polymers also are likely to change the rheology of the air-liquid interface, 4) some of the chemicals are already in formulated spray products and there are human exposure data.

In the following section we have included known results from *in vitro* tests, animal experiments and human exposures. Directly comparing these results to pulmonary surfactant function inhibition is challenging for many reasons, including but not limited to 1) *in vitro* tests using cell culture measure very different endpoints such as cell death or cytokine release that may not translate well to pulmonary surfactant inhibition, 2) *in vitro* tests on cells are almost exclusively done on static cell cultures, whereas pulmonary surfactant function is measured in a dynamic system, 3) animal experiments performed according to guidelines determine lethal concentration, rather than changes in respiration which would be the endpoint most likely correlated with the present results (for a detailed discussion on this issue see (Da Silva et al., 2021b; Hougaard et al., 2023). 4) information from human exposures are most often reported after accidents where the exposed group is small and the aerosol concentration not measured, or in the case of reactions to pharmaceuticals, these represent a sensitive part of the population. The information from these sources can be used in a weight of evidence approach to risk assessment.

The use of these data in risk assessment is further complicated by the physiological function of the lungs. As the lungs have a structure where the internal diameter becomes smaller with each branching of the bronchioles, they are effective filters. The larger the aerosol is, the less likely it will penetrate the airways and enter the alveoli. Therefore knowing the aerodynamic diameter of the aerosol created during spraying is essential, as it will determine which NAM or combination of NAMs is most appropriate to use. When animal experiments are performed according to test guidelines, the aerosol created has to be small enough for animals to inhale. This may not reflect the actual situation when using the product, e.g. one of the tested polymers, BE PVM/MA, has an average particle size that is larger than what is respirable when used (Burnett et al., 2011). Similarly, the packaging of the spray product can affect the size of the aerosols created, e.g. trigger spray vs. pressurized sprays, the geometry of the nozzle and the solvent used in the product. However, in this work we have nebulized all the test chemicals in the same way, and we have not evaluated if real-life products would create aerosols that would reach the pulmonary surfactant layer.

### 4.2 Surfactants

We tested the anionic surfactant SDS as a model for surfactants. SDS is used in cosmetic products and cleaning agents, however it is seldom used in spray products. A mapping of spray cleaning products in Denmark found no products containing SDS, however the “consumer product information database” (https://www.whatsinproducts.com/) has several spray products containing SDS sold in the USA. At high concentrations, SDS is known to cause acute respiratory toxicity, skin and respiratory irritation and therefore routinely used as an *in vitro* positive control in various acute irritation tests (Singer and Tjeerdema, 1993; Welch et al., 2021). We tested SDS at three different aerosolisation rates and five concentrations, increasing the dose-rate led to more severe changes in the dilational rheology (Fig 2a-i). When the normalised change in the complex modulus was plotted against dose-rate, the POD was determined to be 1.63 mg/min, almost 30 times higher than the POD for POTS. SDS was dissolved in water, and water alone did not change the dilational rheology of the cycling pulmonary surfactant, suggesting that the effects are due to the interaction between the aerosolised SDS and the pulmonary surfactant film, rather than the water. SDS was previously found to be a respiratory irritant in mice and guinea pigs (Ciuchta and Dodd, 1978; Zelenak et al., 1982). Dose-dependent toxicity of SDS was observed when tested *in vitro* in a human derived ciliated epithelial cell model (Welch et al., 2021), and in A549 cells (Ritter et al., 2018).

We tested two surfactants classified as quaternary ammonium compounds (QACs); BAC and HTA. QACs are commonly found in spray products, e.g. they were found in 12% of the spray cleaning products on the Danish market (Clausen et al., 2020). The inhibition of pulmonary surfactant function was dose-rate dependent for both, however, BAC had a POD of 0.04 mg/min whereas for HTA it was 0.20. Larsen et al (Larsen et al., 2012) tested both BAC and HTA in a mouse inhalation bioassay, where the breathing patterns of the animals were monitored during exposure. They found that inhalation of QACs led to a reduction in tidal volume, accompanied by an increase in respiratory rate, these changes were attributed to pulmonary irritation. Both HTA and BAC gave rise to dose-rate dependent effects on the reduction in tidal volume, with an increasing concentration in the feed solution, there was a larger depression in the tidal volume. BAC was at least 10 times more potent than HTA at reducing the tidal volume. When we tested these two chemicals for pulmonary surfactant function inhibition, we saw the same relative magnitude in potency, and BAC had a POD slightly lower than POTS.

BAC is a cationic surfactant used in nasal and oral inhalation solutions, household sprays as a disinfectant and a preservative in nasal decongestants (Graf, 1999; Johnson, 2018). In sensitive individuals inhalation of BAC has shown to give rise to respiratory effects. When asthmatic cleaners were challenged with BAC this resulted in effects such as reduced respiratory flow rate and bronchoconstriction (Clausen et al., 2020), and in rare cases in patients after repeated use of BAC-containing nebulizer therapy. Clausen et al (Clausen et al., 2020) concluded that the assessment of respiratory effects in humans after QAC inhalation is difficult because of the uncertainty in exposure, presence of other chemicals and lack of understanding of mechanism of action. When tested on rats, BAC induced a strong inflammatory response in the lungs (Swiercz et al., 2008; Swiercz et al., 2013). It also induced cytotoxicity in A549 cells (Jeon et al., 2019; Kanno et al., 2020; Kim et al., 2020). Furthermore, a dose-dependent increase in surface pressure of the Langmuir-Blodgett pulmonary surfactant monolayer was observed in the presence of BAC (Kanno et al., 2020). However, due to the high droplet size generated from cleaning trigger sprays (median aerosol diameters range from about 70 µm up to well over 100 µm) (Delmaar and Bremmer, 2009) it is unlikely that BAC droplets would reach the alveolar region and produce these effects at normal use levels.

The final surfactant we tested was TX-100, a non-ionic surfactant that is used extensively in the laboratory to induce cell death *in vitro*. TX-100 is classified as a substance of very high concern (SVHC) under REACH because of its endocrine disrupting properties in the environment(European Chemicals Agency, 2012) and is not used in consumer products. TX-100 is very viscous, and dissolving and aerosolising the chemical was limited by this, therefore the highest tested dose-rate was 5 mg/min, and this did not result in a change in the dilational rheology of the pulmonary surfactant film. This dose-rate is 80 times higher than the POD for POTS. TX-100 was tested in a CDS setup by Liu et al (Liu et al., 2024), where it inhibited pulmonary surfactant function by increasing the minimum surfaces tension. However, the difference in experiments performed in the present paper is that the pulmonary surfactant and the test chemicals were mixed prior to testing, rather than aerosolising. When TX-100 was tested on Syrian Hamsters, there were minimal pulmonary pathological changes at the lethal concentration, but shortness of breath was observed within 1-2 h of exposure (Damon et al., 1982). *In vitro*, TX-100 tested on A549 cells induce dose-dependent toxicity (Boesewetter et al., 2006), and in human precision-cut lung slices induced severe tissue damage (Patel et al., 2023).

### 4.3 Polymers

Polymers have so far been exempted from evaluation and registration under REACH, however their regulation is under evaluation. Currently there is no official revision, but preliminary documents suggest that some polymers would require to be registered after the revision (European Chemicals Agency, 2023; Rovida et al., 2023). As mentioned above, even though they were originally exempted from REACH, their safety when used in spray products still need to be determined. As a model for polymer chemicals we tested PHMB, a member of the polymeric guanidine family of chemicals that is widely used as a biocide and preservative in industrial, pharmaceutical and consumer products (Ikeda et al., 1984). We tested PHMB (dissolved in water) at three different aerosolisation rates, and three different concentrations. Similar to SDS, changing the dose-rate of PHMB by either increasing concentration or aerosolisation rate induced a larger change in the dilational rheology. PHMB had a POD of 2.9 mg/min, almost 50 times higher than for POTS.

The antimicrobial activity of PHMB is attributed to the biguanide groups that interact with the polar head groups of the lipids in the bacterial cell membrane, disrupting the organization of the phospholipid bilayer followed by the death of the organism (Ikeda et al., 1984). As pulmonary surfactant is made up of 90% phospholipids, this is likely also the mechanism of toxicity in the inhibition of pulmonary surfactant function. The use of PHMB is not advisable in spray products in the EU(European Union, 2012). When PHMB was tested on A549 cells, this induced a dose-dependent inflammatory response (Kim et al., 2017).

In addition to PHMB we tested five other polymers, BE PVM/MA, PHMG, acrylate co-polymer, polymer 1, and POTS (positive control).

BE PVM/MA has a history of safe use in aerosol hair sprays at concentrations of 2-14% (Burnett et al., 2011), supported by traditional toxicological data. BE PVM/MA dissolved in ethanol inhibited pulmonary surfactant function and a POD of 0.74 mg/min was derived, about 12 times higher than for POTS. Intratracheal installation of a similar copolymer for 13 weeks in rats resulted in an increase in lung weights and increased inflammatory and immune responses with chronic tissue damage in the high dose group (Carthew et al., 2006).

PHMG is a guanidine-based polymer that was used in humidifier disinfectants in Korea due to its bactericidal properties. An epidemiological investigation conducted by the Korean Centre for Disease control and Prevention indicated PHMG as the causative ingredient of the outbreak of fatal lung injury that occurred in pregnant and post-partum women and their children when the chemical was inhaled during its use as humidifier disinfectant (Park et al., 2014; Lee and Yu, 2017). When we tested PHMG for pulmonary surfactant inhibition it did not change the dilational rheology of the pulmonary surfactant film at dose-rates up to 125 mg/min, a dose-rate 2000 times higher than the POD for POTS.

As PHMG was suspected to injure a large group of people it has been extensively studied both *in vitro* and *in vivo*. Inhalation exposure to PHMG aerosols in rats resulted in alveolar emphysema, lung inflammation and pulmonary fibrosis (Song et al., 2014; Yang et al., 2023). Furthermore, irregular and noisy respiration was observed in pregnant rats (Lee et al., 2021) when exposed to PHMG aerosols. Similar pulmonary pathological changes were observed in mice exposed to PHMG aerosols(Song et al., 2018; Li et al., 2021). *In vitro*, PHMG exposure in human derived 3D airway models, EpiAirway™ and MucilAir™, resulted in increased inflammatory and fibrotic responses (Kim et al., 2022). Li 2021 investigated the molecular mechanisms of PHMG induced pulmonary fibrosis, and reported pulmonary surfactant function inhibition as a potential mechanism of toxicity(Li et al., 2021). PHMG was mixed with Curosurf (the same pulmonary surfactant used in present study) an hour before dynamic cycling (3 seconds per cycle) of pulmonary surfactant drop in the CDS. As for other reported results for TX-100, this experiment with PHMG may seem to contradict the results found in Fig 4, however the method of exposing the pulmonary surfactant is different, underscoring the different effects chemicals have on pulmonary surfactant function when the chemicals is mixed with the pulmonary surfactant, or as we have done, exposed to already formed pulmonary surfactant layer. In addition the amount of PHMG compared to the pulmonary surfactant was very high when a change was observed (1:10)(Li et al., 2021).

From our results and the above studies it is likely that the toxic effects seen from inhaling PHMG added as a disinfectant in humidifiers is not from inhibition of pulmonary surfactant function, but rather direct injury to the cells in the lungs or a different mechanism of toxicity. This illustrates the need for more than one NAM to assess the safety of a novel chemical to ensure that the critical toxicological effect is detected.

Acrylate co-polymer has a history of safe use as an addition to hairsprays at concentrations up to 6% (Cosmetic Ingredient Review, 2018). It did not affect the pulmonary surfactant function at dose-rates up to 50 mg/min, a rate 800 times higher than the POD for POTS.

Finally, we tested polymer 1 for its interaction with pulmonary surfactant function *in vitro* as a step in hazard characterisation of a polymer not presently used in spray products. Polymer 1 is a siloxane polymer and it was tested in 2-propanol, it had a POD of 0.20 mg/min, the POD is approximately 3 times higher than POTS. As the POD is very close to that of POTS, a chemical that has known effects on lung function of exposed mice, it would flag a potential safety liability and further information might be needed. For example, a detailed exposure assessment would be required to determine the likely local lung deposition for different product types, to properly characterise the risk. As stated previously, this NAM only covers one of many likely endpoints in the lungs, and further testing would be advisable. We found no relevant literature on the testing of this polymer for effects on the lungs.

### 4.4 Dose-rate dependant toxicity

We have demonstrated that inhibition of pulmonary surfactant function is dependent on the dosing rate of the aerosolised chemical. Comparing this result to alterations in respiration function *in vivo*, either in accidentally exposed humans or in exposed experimental animals, is challenging for several reasons. Firstly, the disruption of pulmonary surfactant function does not have a clear endpoint in intact lungs. In AOP 302 disruption of pulmonary surfactant function is linked to decreased lung function (Da Silva et al., 2021c), but decreased lung function is a broad term. The most convincing results that inhibition of pulmonary surfactant function has a direct effect on the lungs comes from numerous case studies of inhalation of impregnation products. The symptoms range from self-reported and observed symptoms such as coughing, difficulty breathing, tightness in the chest, dyspnoea, nausea and vomiting, to measured effects such as changes in blood samples, spirometry, ground glass opacities on the lungs and infiltrative shadows seen in x-rays, changes seen on the lungs from computerised tomography scans, and reduced SPO_2_. Symptoms occur during or shortly after spraying, and similar symptoms occur when experimental animals are subjected to the same products, but can only be observed if breathing is monitored. The second reason that comparing *in vitro* results to *in vivo* results is challenging is that the acute inhalation studies that are performed according to test guidelines (e.g. OECD TG403, 433 and 436) do not require the breathing of the animals to be monitored during or after exposure, and therefore these symptoms can go unnoticed if only acute inhalation toxicity data is available. Changes in respiration are sometimes noted as clinical signs of exposure, but these are registered cage side, thus have to be severe to be noticed. Thirdly, respiration can change for a variety of reasons in intact lungs, e.g. due to stimulation of nerve endings in the upper or lower parts of the airways, irritation of the airway epithelium or disruption of the air-blood barrier. Studies that link the effect on pulmonary surfactant function with the change in respiration are mostly designed to test the same substances both *in vitro* and *in vivo*, if the tests are only performed *in vivo* these need to monitor breathing during exposure.

In the experiment performed in the present paper we show that the effects on pulmonary surfactant function is dependent upon dose-rate. Most experiments that explore the impact of chemicals or particles on the lungs use a single, high-dose administration method. Therefore, they are not ideal for investigating the effects that depend on the rate at which the dose is given. Baisch et al (Baisch et al., 2014) studied the effects of a bolus of a test material (TiO_2_ nanoparticles) and inhalation of the same dose, the administration of the exposure had different results on lung endpoints, e.g. neutrophil influx. Dose-rate dependent toxicity has been observed for other substances than those studied in this paper, e.g. nanomaterials and substances found in the work environment. Exposure to ZnO, TiO_2_, Al_2_O_3_ and CeO_2_ nanoparticles induced a reduction in tidal volume, accompanied by increase in respiratory rate in a concentration dependent manner. For all nanomaterials, except ZnO, the effect reversed after end of exposure (Larsen et al., 2016). ZnO was speculated to inhibit pulmonary surfactant function and this was later elucidated using the CDS (Larsen et al., 2020), where it was shown that it was the ZnO nanoparticles, rather than Zn-ions that inhibited pulmonary surfactant function *in vitro*. Dose-rate dependent respiratory effects have also been shown for butter flavouring agent diacetyl (2,3-butanedione) known to cause bronchiolitis obliterans in exposed workers (Larsen et al., 2009). Mice exposed to the substance show a concentration dependent reduction in tidal volume and breathing rate, the substance also caused sensory and pulmonary irritation.

In previous studies of the inhibition of pulmonary surfactant (Beck-Broichsitter et al., 2017; Kondej and Sosnowski, 2019; Da Silva et al., 2021a; Van Bavel et al., 2023), microscopy found that the structures which form during compression are altered by the presence of a test chemical. It was observed by (Van Bavel et al., 2023) that vaping additives impaired the formation of multilayer structures. Similar alterations to the microstructure were seen in cryo-TEM experiments wherein pulmonary surfactants were exposed to three chemicals known to show acute inhalation toxicity (Da Silva et al., 2021a). These structures are linked to the increased elasticity of pulmonary surfactant monolayers as well as preserving the structural integrity of the monolayer at extreme compression (Alonso et al., 2004; Liekkinen et al., 2020; Castillo-Sanchez et al., 2021). This therefore suggests that the inhibition of the function of pulmonary surfactant may be related to hydrophobic interactions with SP-B and SP-C proteins. These interactions then inhibit the formation of three dimensional structures and reservoirs. Da Silva et al (Da Silva et al., 2021a) showed film collapse in surface area-pressure isotherms at low surface area per molecule when the pulmonary surfactant is mixed with a test chemical, as opposed to pure pulmonary surfactant samples which show a surface pressure plateau at low surface area per molecule. The mechanism of action of the positive control used in the present work was studied *in vitro* using Langmuir isotherms, it was found that adding SP-B restored the function of the pulmonary surfactant, whereas adding SP-C did not restore function(Larsen et al., 2014).

In our mechanistic description of the inhibition of pulmonary surfactant we assume that interactions between the molecules results in increased rate desorption of the pulmonary surfactant molecules into the bulk from the interface. What is not immediately clear is whether these interactions are dominated by the test chemical interacting with the phospholipid component of the pulmonary surfactant or the protein component. Further experiments could possibly demonstrate this by altering the ratio of phospholipid and protein in a model pulmonary surfactant solution as done in(Larsen et al., 2014).

We tested if the different test chemicals had surface activity as described in the M&M section, five chemicals could not be tested as they did not dissolve in water (BE PVM/MA, HTA, polymer 1, POTS and SRS). We theorized that chemicals with surface activity would inhibit pulmonary surfactant function by competing with pulmonary surfactant at the air-liquid interface. However, of the surfactants, the most surface active was TX-100 (30.8 mN/m at 0.5% w/v) and this did not inhibit pulmonary surfactant function. Of the polymers we could measure surface activity for PHMG, PHMB and Acrylate co-polymer, the first two did not lower surface tension, but Acrylate co-polymer lowered it to 38.5 mN/m and was surface active. However, PHMB inhibited pulmonary surfactant function, whereas PHMG and Acrylate co-polymer did not. We tried to find a physical-chemical characteristic that could divide chemicals that inhibited pulmonary surfactant function from those that did not affect function, however none of the characteristics could do this.

### 4.5 Limitations of the study and future work

This study has several limitations, firstly it is not an “animal-free” method as we use Curosurf as the model surfactant. Curosurf is isolated from pig lungs, and sold as a pharmaceutical. We hope that in the future we can replace this animal-derived component of the test system with a synthetic pulmonary surfactant, however this is not available at the moment. Second, we use the POD of POTS, which is known to cause dose-rate dependent lung toxicity in exposed mice and is restricted for use in spray products in REACH, to hazard rank the tested chemicals. To make a direct comparison between the obtained results to results from exposed humans, animals or cell experiments we would need to determine the inhibitory dose. This is not possible at the moment, however we are working on integrating mass spectrometric measure of the test chemical in the exposed pulmonary surfactant. However, when we have measured and estimated the inhibitory dose in the past, there has, in most cases, not been a direct correlation between what is measured *in vitro* and the estimated deposited dose when an effect on respiration is observed. This is likely because the *in vitro* system contains a finite amount of pulmonary surfactant, whereas the lungs have a more dynamic pulmonary surfactant pool, and other physiological systems that contribute to the effect on breathing.

Inhalation is a major route of human exposure, but the lungs are the organ that has proven as one of the most difficult to replace by *in vitro* methods. This is both because of the anatomy of the lungs with different architecture and cell composition throughout the path of inhaled air, as well as the constant motion of the organ with each breath. Several cell models of different parts of the lungs have been developed, and these cover many endpoints, but none have so far reproduced the function of the essential component pulmonary surfactant. Measuring pulmonary surfactant function *in vitro* fills an important endpoint in evaluating the effect of inhaled chemicals on the alveoli, without testing on experimental animals. It can thus reduce the need for data from animal experiments and in combination with other *in vitro* methods likely replace the reliance on animals for testing portal of entry effects of inhaled chemicals.

As mentioned previously the method does not take into account the formulation of the product, and different precautions that could be taken to minimise the exposure, e.g. by increasing aerosol size to limit inhalation, formulating the product in less volatile solvents to avoid solvent evaporation during application, as this would avoid aerosols shrinking and more aerosols would settle by gravity. Modelling the aerosol deposition in the airways would inform the choice of NAM to use in hazard assessment, however this has not been included in the present work, and would be important in an IATA decision making context. Future work could be focused on comparing the PODs obtained from this study with predictions of human exposure for chemicals for which an exposure scenario is possible to establish, using CFD and MPPD modelling (Ramanarayanan et al., 2022).

In consumer products surfactants are often found as a mixture as they together improve the function of the product (Lindman and Nylander, 2017) and have synergistic effects (Nakama, 2017). If mixtures of different surfactants, or surfactants and polymers can also increase the ability to inhibit pulmonary surfactant function is a task for further exploration.

The study was ground-breaking in establishing PODs that serve as a benchmark for hazard ranking of test chemicals. While BAC was found to induce similar respiratory changes as POTS in mice, albeit at a lower aerosol concentration, it is important to note that BAC has a history of safe use in trigger sprays. The other chemicals tested exhibited PODs ranging from 3 to 80 times higher than that of POTS. Although comparing PODs can provide a relative measure of hazard, it does not offer a definitive assessment of risk without considering the exposure levels associated with product use. For instance, HTA, with a POD about fourfold higher than POTS, elicited effects in experimental animals akin to those observed with both BAC and POTS. Therefore, a comprehensive risk assessment necessitates an evaluation of both the inherent hazard and the potential exposure to humans.

## 5. Conclusion

Our study has examined multiple aspects of the inhibition of pulmonary surfactant by aerosolised chemicals. We have incorporated *in vitro* experiments, rheometric analysis and theory to explore the toxicity of a variety of test chemicals. Our findings suggest that, in agreement with data from exposed humans and animal studies, the inhibition of the pulmonary surfactant function is dependent on the rate of exposure. This is seen for all test chemicals which affected surfactant function in this study.

This study was motivated by the aim to develop an understanding of pulmonary surfactant toxicity using *in vitro* experiments studied previously in animal studies. We have been able to replicate the *in vivo* characteristics and to isolate key aspects of the inhibition, corroborating the findings of both medical and *in vivo* studies. Our work here therefore acts as an encouraging example of how animal testing may be superseded by *in vitro* non-animal methods.

## Acknowledgments

We would like to thank Rikke Nilsson and Karen Bo Frydendall for help with literature search strategy and reference retrieval. Unilever funded the study, JBS and SRS acknowledges funding from FFIKA, Focused Research Effort on Chemicals in the Working Environment, from the Danish Government. HJB and MTS would like to acknowledge Joe Reynolds, Sophie Cable, Anthony Bowden and Iris Muller for useful discussions.

## Data availability

Data is available upon request from.

## Supplementary material

**Supplementary table 1:**
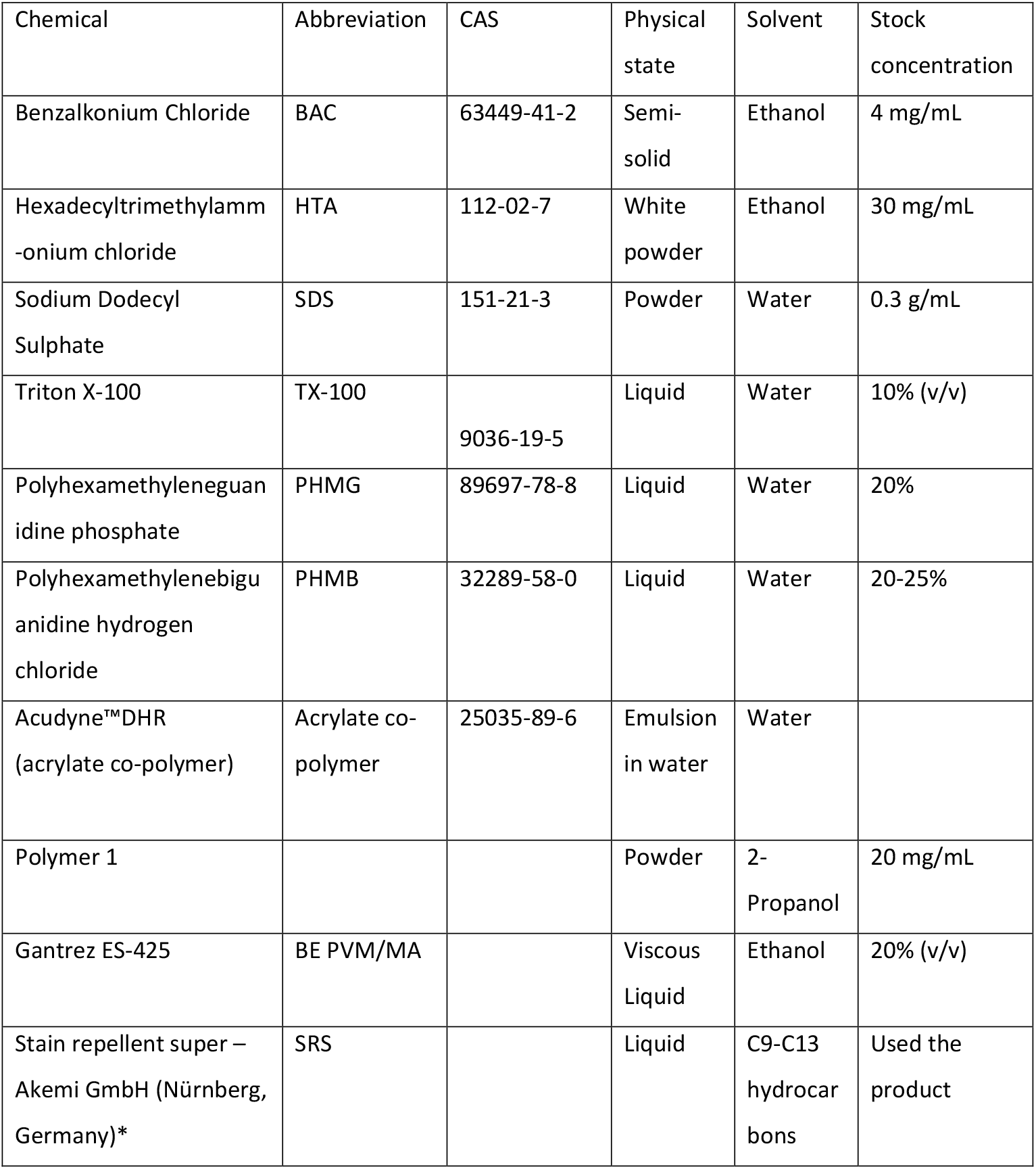
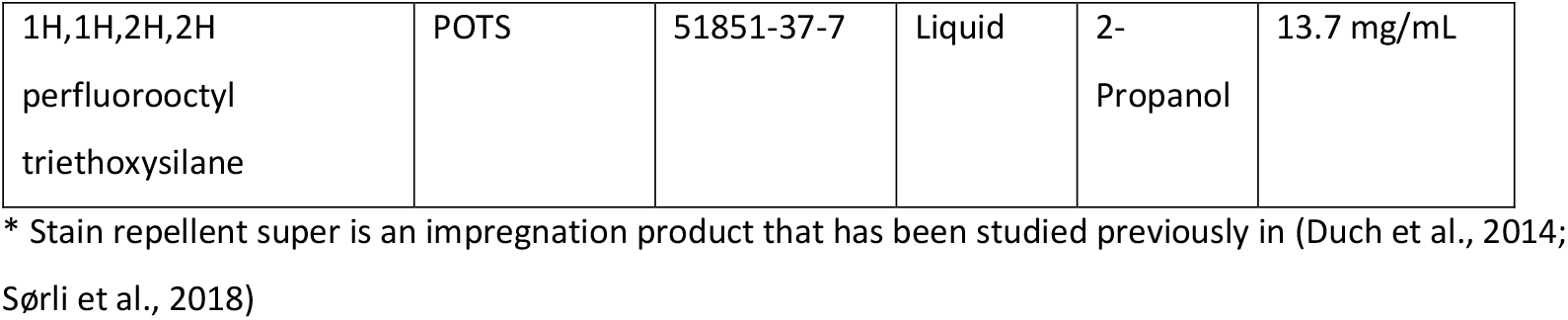
Stock concentrations, solvent and physical state of surfactants and polymers that were tested in the constrained drop surfactometer.

| Chemical | Abbreviation | CAS | Physical state | Solvent | Stock concentration |
| --- | --- | --- | --- | --- | --- |
| Benzalkonium Chloride | BAC | 63449-41-2 | Semi-solid | Ethanol | 4 mg/mL |
| Hexadecyltrimethylamm<br>-onium chloride | HTA | 112-02-7 | White powder | Ethanol | 30 mg/mL |
| Sodium Dodecyl Sulphate | SDS | 151-21-3 | Powder | Water | 0.3 g/mL |
| Triton X-100 | TX-100 | 9036-19-5 | Liquid | Water | 10% (v/v) |
| Polyhexamethyleneguan<br>idine phosphate | PHMG | 89697-78-8 | Liquid | Water | 20% |
| Polyhexamethylenebigu<br>anidine hydrogen chloride | PHMB | 32289-58-0 | Liquid | Water | 20-25% |
| Acudyne™DHR<br>(acrylate co-polymer) | Acrylate co-polymer | 25035-89-6 | Emulsion in water | Water |  |
| Polymer 1 |  |  | Powder | 2-Propanol | 20 mg/mL |
| Gantrez ES-425 | BE PVM/MA |  | Viscous Liquid | Ethanol | 20% (v/v) |
| Stain repellent super –<br>Akemi GmbH (Nürnberg,<br>Germany)* | SRS |  | Liquid | C9-C13 hydrocarbons | Used the product |
| 1H,1H,2H,2H<br>perfluorooctyl<br>triethoxysilane | POTS | 51851-37-7 | Liquid | 2-Propanol | 13.7 mg/mL |

\* Stain repellent super is an impregnation product that has been studied previously in (Duch et al., 2014; Sørli et al., 2018)

### Literature search

For each chemical the following review of the literature was done: 1) we identified if the chemical had a dossier in ECHA 2) a literature search string was used in PubMed (https://pubmed.ncbi.nlm.nih.gov/): search #1 ‘Chemical name’ OR ’CAS number’, #2 ‘pulmonary surfactant’, search #3 ‘inhalation’, search #4 ‘lung’. Search #1 was combined with #2, #3 and #4 consecutively, and the retrieved papers were collected. The list of papers was curated by the relevance of the title, and then by abstract. The number of papers found before and after curation can be found in supplementary table 2. All relevant papers were reviewed and scored using the ‘ToxRTool’ (Schneider et al., 2009), a software-based tool to evaluate the reliability of the data.

**Supplementary table 2:**
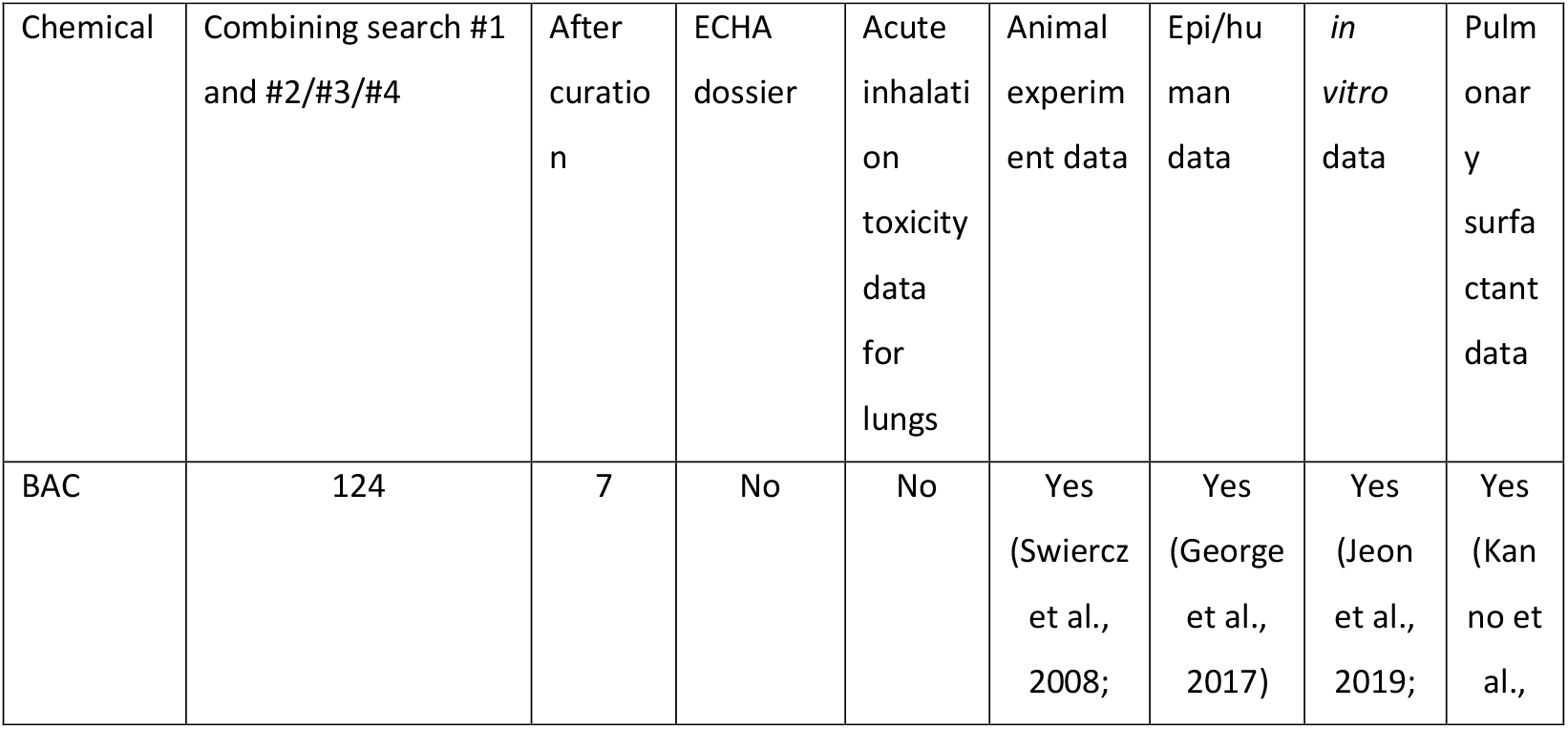
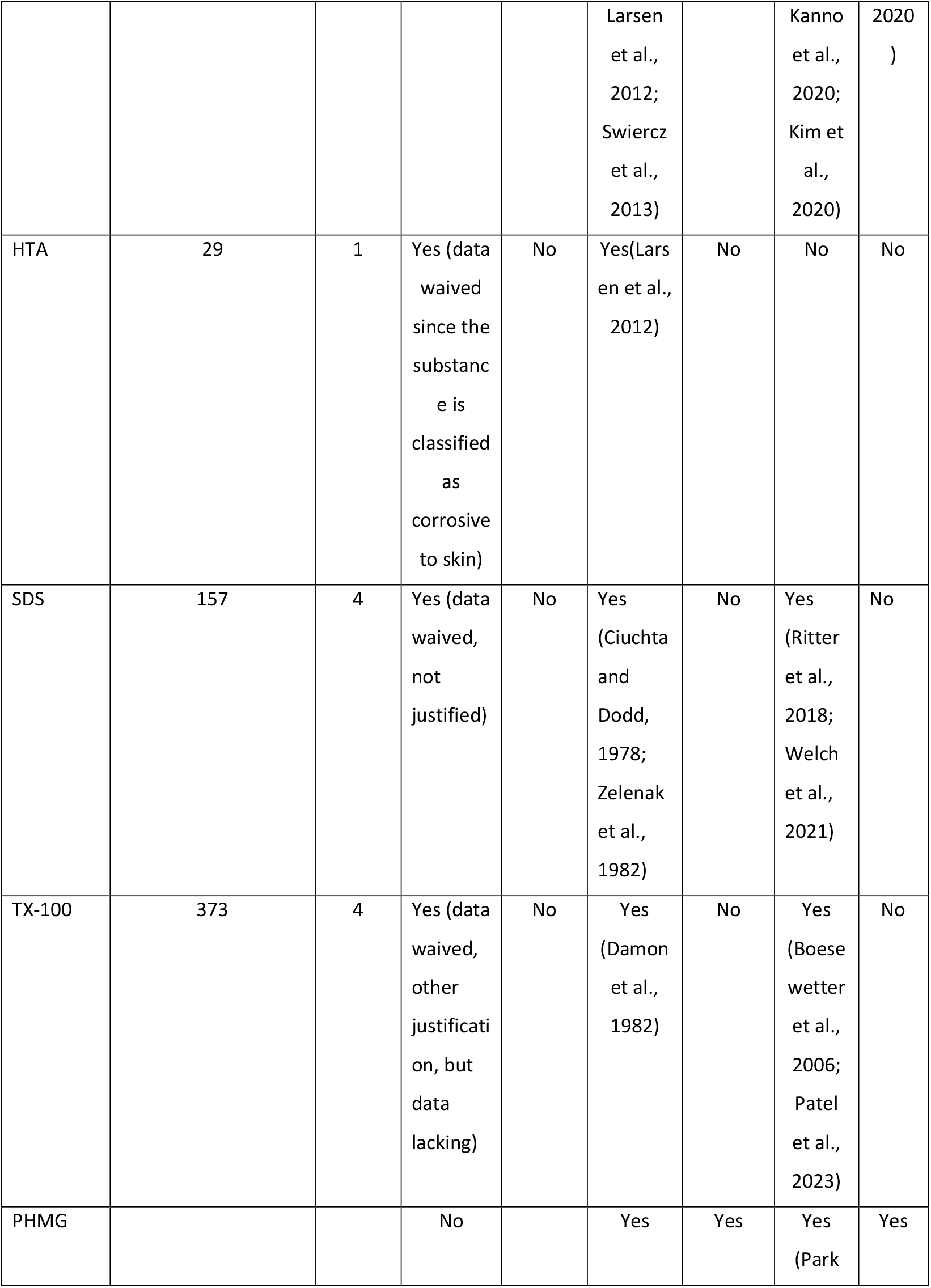
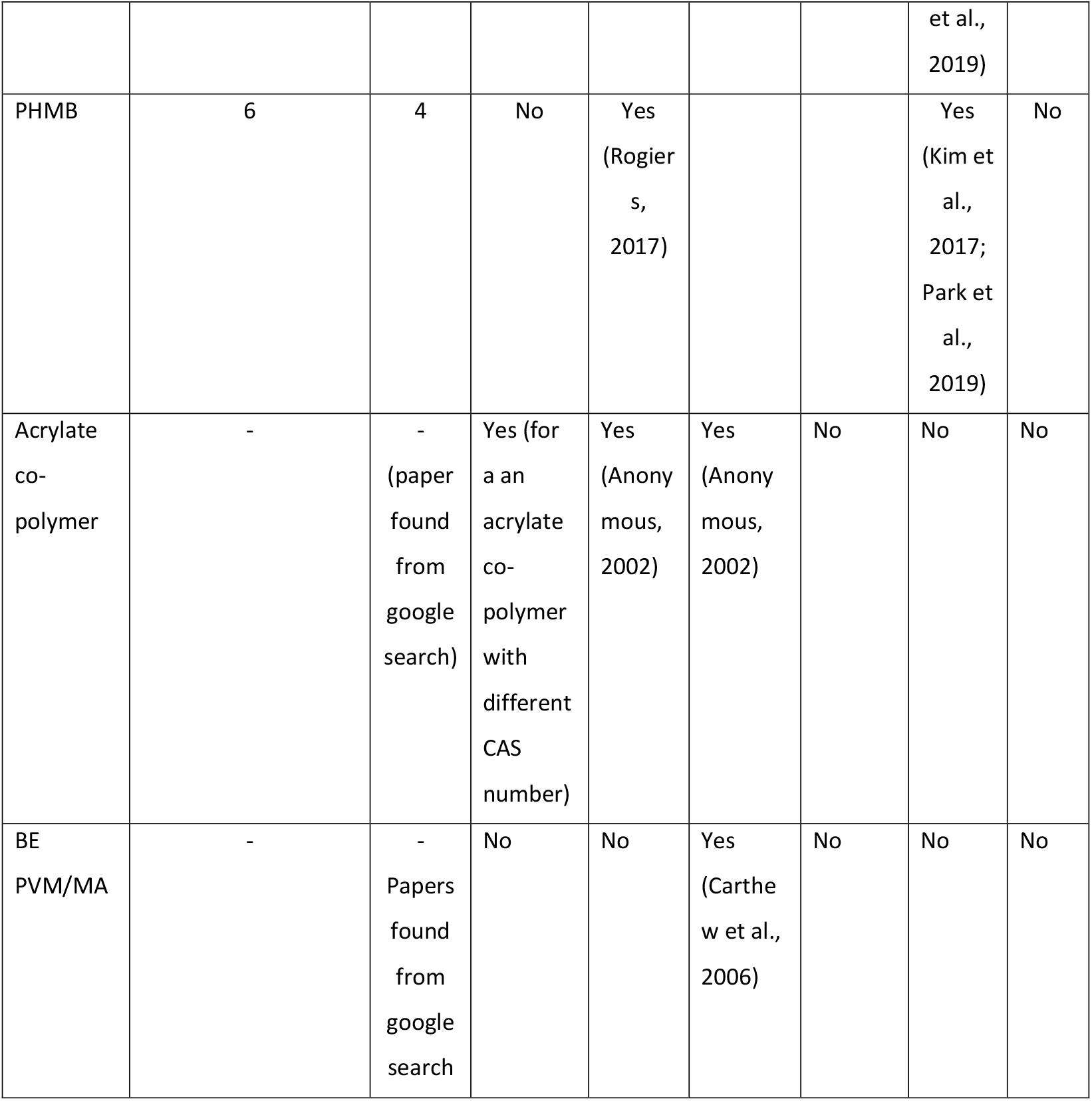
Summary of the literature search conducted in PubMed.

| Chemical | Combining search #1<br>and #2/#3/#4 | After<br>curatio<br>n | ECHA<br>dossier | Acute<br>inhalati<br>on<br>toxicity<br>data<br>for<br>lungs | Animal<br>experim<br>ent data | Epi/hu<br>man<br>data | *in*<br>*vitro*<br>data | Pulm<br>onar<br>y<br>surfa<br>ctant<br>data |
| --- | --- | --- | --- | --- | --- | --- | --- | --- |
| BAC | 124 | 7 | No | No | Yes<br>(Swiercz<br>et al.,<br>2008; | Yes<br>(George<br>et al.,<br>2017) | Yes<br>(Jeon<br>et al.,<br>2019; | Yes<br>(Kan<br>no et<br>al., |
|  |  |  |  |  | Larsen et al., 2012; Swiercz et al., 2013) |  | Kanno et al., 2020; Kim et al., 2020) | 2020 ) |
| HTA | 29 | 1 | Yes (data waived since the substance is classified as corrosive to skin) | No | Yes(Larsen et al., 2012) | No | No | No |
| SDS | 157 | 4 | Yes (data waived, not justified) | No | Yes (Ciuchta and Dodd, 1978; Zelenak et al., 1982) | No | Yes (Ritter et al., 2018; Welch et al., 2021) | No |
| TX-100 | 373 | 4 | Yes (data waived, other justification, but data lacking) | No | Yes (Damon et al., 1982) | No | Yes (Boesewetter et al., 2006; Patel et al., 2023) | No |
| PHMG |  |  | No |  | Yes | Yes | Yes (Park et al.,<br>2019) | Yes |
| PHMB | 6 | 4 | No | Yes<br>(Rogier<br>s,<br>2017) |  |  | Yes<br>(Kim et<br>al.,<br>2017;<br>Park et<br>al.,<br>2019) | No |
| Acrylate<br>co-polymer | - | -<br>(paper<br>found<br>from<br>google<br>search) | Yes (for<br>a an<br>acrylate<br>co-polymer<br>with<br>different<br>CAS<br>number) | Yes<br>(Anony<br>mous,<br>2002) | Yes<br>(Anony<br>mous,<br>2002) | No | No | No |
| BE<br>PVM/MA | - | -<br>Papers<br>found<br>from<br>google<br>search | No | No | Yes<br>(Carthe<br>w et al.,<br>2006) | No | No | No |

